# Longitudinal study of the udder microbiome of Norwegian Red dairy cows using metataxonomic and shotgun metagenomic approaches: Insights into pathogen-driven microbial adaptation and succession

**DOI:** 10.1101/2025.04.04.647183

**Authors:** Vinicius da Silva Duarte, Fiona Valerie Franklin, Alicja Krysmann, Davide Porcellato

## Abstract

Bovine mastitis remains the most significant disease affecting dairy herds globally, driven by its multi-etiological nature and the complex dynamics of udder immunity and infection. While research addressing the microbial and immunological aspects of the bovine udder is limited, optimizing the udder microbiome has emerged as a promising strategy for preventing mastitis. This longitudinal study aimed to investigate the udder microbiome throughout lactation using both metataxonomic and shotgun metagenomic approaches, including analysis at the metagenome-assembled genome (MAG) level. The use of such methodologies provides a deeper understanding of the microbial composition and dynamics within the udder, bridging critical gaps in knowledge and revealing potential interactions between the microbiota and host. Milk samples were collected from 342 individual quarters of 24 Norwegian Red dairy cows. Significant variations in somatic cell count and microbiota composition were observed across lactation stages. Quarters with low somatic cell count were notably higher during early lactation (80%) and mid-lactation (78.9%) compared to dry-off (53.1%) and late lactation (53%), with high somatic cell countobserved in 20–47% of samples. Diversity indices based on Shannon and Simpson metrics indicated significant effects of lactation stage, somatic cell count, and individual animal variability on microbial diversity. PERMANOVA analyses confirmed that individual animal variability (15.73%) and lactation period (5.52%) were the strongest factors influencing microbiota structure, with dysbiosis linked to mastitis-causing pathogens contributing 7.17% of the variance. Key pathogens identified included *Enterococcus faecalis*, *Staphylococcus aureus*, *Streptococcus uberis*, and *Staphylococcus chromogenes*, with persistent infections observed for *S. aureus* and *S. chromogenes*, but not *S. uberis*. Samples with low somatic cell count were enriched in beneficial genera such as *Corynebacterium*, *Bradyrhizobium*, and *Lactococcus*, while *Staphylococcus* predominated in milk samples with high somatic cell count. Dimensionality reduction integration with culturomics enhanced milk microbiota classification, providing novel insights into udder microbiota dynamics and their role in bovine mastitis. Metagenomic shotgun sequencing revealed pathogen-specific metabolic signatures in the bovine mammary gland, identifying 289 MetaCyc pathways. Genome-centric analysis reconstructed 142 metagenome-assembled genomes, including 26 from co-assembly and 116 from individual assembly. Multi-locus sequence typing, virulence factors, and antimicrobial resistance gene profiling provided insights into pathogen adaptation and persistence in the bovine mammary gland. Notably, 27 bacteriocin gene clusters and 322 biosynthetic gene clusters were predicted using genome mining tools. Our findings contribute to the establishment of pathogen-based therapies and enhance our understanding of the milk microbiome, which remains far from fully characterized. Such insights are vital for developing effective strategies to combat mastitis and improve dairy cattle health.

## 1. Introduction

Bovine mastitis remains the leading disease affecting dairy herds worldwide ^1^. Due to its multi-etiological nature, it is a challenging condition to eradicate and is primarily managed through husbandry practices and the use of antibiotics to treat bacterial infections ^2^, which are the primary cause of intramammary infections (IMIs).

Mastitis can be classified into clinical and subclinical forms based on clinical features. Clinical mastitis (CM) is marked by distinct symptoms, such as the presence of flakes, clots, or watery secretions in milk. Affected quarters typically show swelling, warmth, and pain. In acute cases, systemic signs like hyperthermia, anorexia, and depression may also occur. The consequences of CM can be severe, often leading to cow mortality, agalactia, or premature culling ^3^. Subclinical mastitis (SCM), on the other hand, is primarily identified by an increased somatic cell count (SCC) and reduced milk production, with SCC being considered the diagnostic gold standard for detecting SCM ^4^.

Microbiological culture of milk remains the gold standard for pathogen identification, despite some limitations ^5^. Few studies have investigated the microbial composition of milk samples classified as “mixed growth,” which are often considered contaminated during sampling ^6^, culture-negative samples from clinical mastitis ^7^, and samples from cows or quarters with high SCC (subclinically affected quarters) ^8^. Overall, both mixed growth and culture-negative samples constitute a substantial proportion (30% or more) of the milk samples submitted for mastitis diagnostics ^9^.

Recent advancements in high-throughput next-generation sequencing (NGS) technology and bioinformatics tools over the past decade have facilitated a shift from traditional clinical microbiology to ecological and genomic characterization of the microbiome associated with infections in different research fields ^10^, including udder health ^11^. This transition does not aim to replace traditional clinical microbiology, but rather emerges as a complementary diagnostic tool and a means to study the interactions between the host and its microbiome ^12,13^. For example, studies employing metataxonomic analysis have shown that culture-based diagnoses generally align with the most prevalent organisms identified through metagenomic sequencing, although additional potential pathogens have been detected in culture-negative samples ^14^.

The dynamics of udder immunity and infection are complex, influenced by factors such as the cow’s health status, environmental conditions, and the resident microbiota ^15^. Research has identified distinct changes in microbiome composition between healthy and mastitic milk, with key phyla such as *Proteobacteria*, *Bacteroidetes*, *Firmicutes*, and *Actinobacteria* playing significant roles^16^. Metagenomic analysis has further revealed previously unreported opportunistic strains in clinical mastitis samples, along with functional pathways related to bacterial colonization and antibiotic resistance ^17^. However, the etiology of udder dysbiosis remains unclear, particularly whether it serves as a cause or consequence of disturbances in host-specific factors, health status, environmental conditions, xenobiotic exposure, or inadequate hygiene. While comprehensive studies directly addressing this question in bovine udders are limited, research on lactating mothers can provide valuable insights ^18^. Additionally, as reported by Urrutia-Angulo et al. (2024)^19^, the clinical definition of udder health status does not consistently align with the microbial profile.

Evidence suggests that SCM may have a distinct pathological origin compared to CM, with unique microbial signatures and sensory protein profiles ^20^. These findings highlight the significant potential of metagenomic approaches to improve mastitis diagnosis and advance our understanding of its pathophysiology. Metagenomic studies provide valuable insights into the temporal dynamics of microbial communities and ^21^, when paired with metagenome reconstruction, reveal possible interactions within the microbiota (ecological studies) and between the microbiota and the host ^22,23^. Furthermore, when integrated with community proteogenomics, these approaches become instrumental in identifying potential biomarkers for the early diagnosis of subclinical mastitis and in elucidating the pathophysiology of both subclinical and clinical mastitis, which remains incompletely understood ^24,25^.

Optimizing the udder microbiome has been proposed as a promising strategy for preventing mastitis in dairy cattle, as a healthy microbiome contributes to protection against pathogen colonization and the overgrowth of opportunistic pathobionts ^26,27^. Therefore, this longitudinal study aimed to investigate the udder microbiome throughout lactation using metataxonomic and shotgun sequencing approaches. Studies of this nature contribute to establishing pathogen-based therapies and provide insights into the composition of the milk microbiome, which remains far from fully characterized. This gap in knowledge motivated the use of shotgun metagenomics and analysis at the metagenome-assembled genome (MAG) level in this study since such comprehensive investigations remain relatively scarce.

## 2. Material and methods

### 2.1. Experimental design and sample collection

Twenty-four Norwegian Red cows were chosen from the “Centre for Livestock Production” at the Norwegian University of Life Sciences. The farm adheres to the regulations set by the Norwegian Food Safety Authority for food production and animal welfare. Sample collection and the use of related information were approved by the farm owners, and no invasive procedures were involved in the study. The dairy cows were kept in freestalls equipped with rubber mats and raw wood chips for bedding. Their diet included silage available at all times and pelleted feed, adjusted according to each cow’s milk production.

The experimental period took place between the autumns of 2022 to 2023. Milk samples were collected at four different time points, covering two lactation cycles: i) before dry-off in 2022 (DO-2022), ii) early lactation (EL-2023), iii) mid-lactation (ML-2023), and iv) late lactation (LL-2023). On each sampling day, hindmilk was aseptically collected from each quarter at the end of the regular milking routine, as described by Porcellato et al. (2020) ^28^. In short, after the milking apparatus was removed, the teats were cleaned with iodine and then 70% ethanol, and approximately 350 mL of milk was manually collected. The procedure followed the guidelines of the National Mastitis Council (NMC; www.nmconline.org). Once collected, the milk samples were transported on ice to the laboratory and then divided into polystyrene tubes. Fresh milk samples were used for microbiological analyses (item 2.2), while milk samples used for metagenomic analyses were immediately frozen at -20°C until further processed (items 2.3 and 2.4).

### 2.2. BacSomatic, culturing and identification of isolates

Milk samples from all quarters were sent to the TINE Mastitis Laboratory in Molde, Norway, for microbiological analysis and species identification (gold standard diagnostic). Bacteriological culturing followed standard protocols 2.3. In summary, 10 μL of milk was plated on cattle blood agar with esculin and incubated at 37°C. The plates were examined after 24 and 48 hours. Species identification was then carried out using MALDI-Tof MS (Microflex LT system. Bruker Daltonics).

Concomitantly, the same fresh milk samples were also plated in four different media at SciFood (KBM. NMBU) and incubated at 37°C for 24, 48 and 72 hours as follows: Tryptic Soy Agar (TSA, Sigma-Aldrich) plus 0.01% Tween 20, Blood agar (Thermo Scientific), brilliance CRE agar (detection of *Enterobacteriaceae* Producing KPC-and Metallo-Carbapenemase, ThermoFisher Scientific) and brilliance ESBL agar (detection of extended-spectrum beta-lactamase-producing *Enterobacteriaceae*, Thermo Scientific). Lastly, the samples were analyzed for somatic cell count (SCC) and individual bacterial count (IBC) using the BacSomatic instrument (Foss Electric, Hillerød, Denmark). In this study, samples were classified as either negative or positive for mastitis-causing pathogens based on the official mastitis report.

### 2.3. Metagenomic DNA extraction and 16S rRNA gene amplicon sequencing

#### 2.3.1. Bacterial pellet preparation and metagenomic DNA extraction with DNeasy PowerFood Microbial Kit

For metagenomic DNA (mgDNA) extraction, a bacterial pellet was isolated from 40 mL of milk as outlined by Winther et al. (2022) ^21^. A total of 345 milk samples were processed. Initially, 40 mL of milk was thawed overnight at 4°C and centrifuged at 8,000 × g for 10 minutes at 4°C. The whey fraction was discarded, and the bacterial pellet along with the fat layer was washed on an orbital shaker at 250 rpm for 15 minutes using a solution of 2% citrate water and 0.1% Tween 20. The mixture was then centrifuged again at 8,000 × g for 10 minutes at 4°C, after which the supernatant was removed, and the fat layer was gently collected and after discarded using a sterile cotton swab soaked in 70% ethanol. The bacterial pellet was then harvested and washed with 2% citrate water (16,200 × g for 3 minutes).

The metagenomic DNA (mgDNA) was extracted using the DNeasy PowerFood Microbial Kit (Qiagen. Düsseldorf. Germany). The bacterial pellet was transferred directly into a PowerBead tube and placed in a FastPrep-24 5G Instrument (MP Biomedicals European HQ) for three rounds of bead-beating at (*Lactococcus* program, 30 seconds with a 5-minute cooling period between each round). The samples were then processed following the manufacturer’s instructions. Finally, the mgDNA was eluted in 50 µL of elution buffer and stored at −20°C until further processed.

#### 2.3.2. Library preparation and 16S rRNA gene amplicon sequencing

Library preparation for amplicon sequencing followed the method previously described in section 2.3.1. Specifically, the V3 and V4 regions of the 16S rRNA genes were amplified using the primers Uni340F (CCTACGGGRBGCASCAG) and Bac806R (GGACTACYVGGGTATCTAAT). The PCR reagents and amplification conditions matched those outlined by Porcellato et al. (2020)^28^. Negative controls consisting of only reagents and nuclease free water were included to check for contamination during DNA extraction and library preparation. Sub-libraries were cleaned and normalized using the SequalPrep Normalization Plate (96) Kit (Thermo Fisher Scientific. USA) and then pooled. The final library concentration was measured with Qubit 2 using the dsDNA HS kit (Thermo Fisher Scientific. USA), and sequencing was performed on an Illumina NovaSeq 6000 platform (Illumina) using the 2 × 250 bp V3 kit (Illumina) at Novogene (Cambridge. UK). A total of four sequencing batches were completed.

### 2.4. Metagenomic DNA extraction and shotgun metagenomic sequencing

#### 2.4.1. Metagenomic DNA extraction with MolYsis complete5 kit

For metagenomic shotgun sequencing, 73 samples were selected based on their SCC levels and their representation of all the major mastitis-causing pathogens identified among the samples included in the study. The bacterial pellet was obtained as outlined in section 2.3.1 and processed using the manufacturer’s protocol (Molzym GmBH & Co. KG. Bremen. Germany). In brief, cells were resuspended in 1 mL of buffer SU, and host cells were lysed by adding 250 µL of chaotropic buffer (buffer CM). The released nucleic acids were degraded using an enzyme (MolDNase B). Microbial cells were then pelleted and lysed with specific reagents and proteinase K. The mgDNA was isolated using spin columns, and 50 µL of DNA was eluted and stored at −20°C for further processing.

#### 2.4.2. Multiple displacement amplification by Phi29 DNA polymerase

Following the manufacturer’s instructions, whole metagenome amplification was conducted on 73 samples using multiple displacement amplification (MDA) with the REPLI-g Single Cell kit (Qiagen: 150345). Briefly, 5 µL of metagenomic DNA (mgDNA) was added to a microcentrifuge tube along with 5 µL of buffer D1 and mixed by vortexing. After a 3-minute incubation at room temperature (15–25°C), 10 µL of buffer N1 (stop solution) was added. To the 20 µL of denatured mgDNA, 30 µL of master mix (comprising 29 µL REPLI-g mini reaction buffer and 1 µL REPLI-g mini DNA polymerase) was added. The final mixture (50 µL) was gently mixed, incubated at 30°C for 16 hours, and then heat-inactivated at 65°C for 3 minutes using the SimpliAmp (Applied Biosystems). The mgDNA concentration was measured using the Qubit HS dsDNA assay Fluorometer (Thermo Fisher Scientific. USA) and stored at −20°C for further analysis.

#### 2.4.3. Library preparation and sequencing

For Illumina sequencing library preparation, the mgDNA concentration of the MDA-treated samples was measured using the Qubit HS dsDNA kit. If the extracted DNA concentration was below 0.5 ng/µL, 5 µL of the undiluted sample was used. Otherwise, the DNA was diluted to 0.2 ng/µL. DNA preparation for sequencing followed the guidelines of the Illumina Nextera XT Library Preparation Kit. The DNA concentration was measured using the Qubit HS dsDNA assay, and the concentration was calculated before being diluted and pooled at equimolar ratios. The DNA library was sequenced on the Illumina NovaSeq 6000 system (2 × 150 bp read length) at Novogene (Cambridge. UK).

### 2.5. Bioinformatics processing data

#### 2.5.1. 16S rRNA gene amplicon sequencing

The demultiplexed raw paired-end reads obtained from sequencing were uploaded and processed in QIIME2 (version 2021.4) using the Casava 1.8 paired-end pipeline ^31^. DADA2 was selected for its ability to enhance taxonomic resolution through precise identification and error correction of sample sequences that differ by as little as a single nucleotide. This process involved several sequential steps, including filtering, trimming, denoising, dereplicating, merging paired reads, and removing chimeric sequences ^32^. Subsequently, amplicon sequence variants were used to generate a phylogenetic tree through the align-to-tree-mafft-fasttree pipeline in the q2-phylogeny plugin ^33^. Taxonomy assignment for the 16S data was performed using a Naïve Bayes pre-trained Silva-138–99-nb-classifier ^34^.

For downstream metataxonomic analysis, QIIME artifacts were imported into R (R Core Team 3.6.2. 2019) using the qiime2R package v.099.20 (https://github.com/jbisanz/qiime2R; accessed on 5 November 2024). *In silico* contaminant identification and removal of amplicon sequence variants (ASVs) were conducted for each sequencing batch using the R package Decontam (version 1.12) ^35^, employing the frequency method with a user-defined threshold of 0.5. After decontaminating the samples with the help of negative controls and before calculating any diversity indices, ASVs assigned to *g-Mitochondria*. *o-Chloroplast*, and *d-Archaea*, assigned as Unknown at the phylum level, were excluded from the dataset. To improve the taxonomical assignment and make inferences at the species level when appropriate, ASVs were manually verified using the MegaBLAST search against the 16S ribosomal RNA sequences (Bacteria and Archaea) database.

#### 2.5.2. Shotgun sequencing

Short reads from the metagenomic datasets were processed using the metaWRAP v1.3^36^ pipeline, following the bioinformatics workflow for genome-centric metagenomics of the bovine hindmilk microbiome as previously outlined by Duarte & Porcellato (2024)^23^. In short, the quality of the raw sequence data was verified using the FastQC tool (https://www.bioinformatics.babraham.ac.uk/projects/fastqc/). The raw reads were trimmed with Trim-galore v0.4.3 ^37^, and then, the bovine-derived reads were removed with bmtagger v3.101 (default settings). The high-quality reads were assembled using both the individual or co-assembly methods with MegaHit v1.0^38^. Subsequently, the assemblies were binned with metaBAT v2.12.1 (-m 1500 and -- unbinned parameters) ^39^, Maxbin v2.2.4 (-markerset 40 option) ^40^ and CONCOCT v0.4.0 (default settings) ^41^ using the metaWRAP-Binning module. The resulting three bin sets were consolidated with metaWRAP-Bin-refinement, and the completion and contamination of the resulting bins were evaluated with CheckM v1.0.7 (default settings) ^42^. Taxonomic and functional profiles were obtained using short reads with MetaPhlAn 3.0 ^43^ and HUMAnN 3.0 ^43^, respectively.

All bioinformatics analyses were performed with the Orion Cluster at NMBU. The raw reads were deposited into the Sequence Read Archive database (http://www.ncbi.nlm.nih.gov/sra) under the BioProject PRJNA950968.

#### 2.5.3. MAG data analysis

Metagenome-assembled genomes (MAGs) were classified into high, medium, and low quality according to the minimum information about metagenome-assembled genome guidelines (MIMAG) ^44^. GTDB-Tk (v2.2.5, reference data version r207_v2) ^45^ was used for taxonomic classification. Identifiers were assigned to the MAGs based on taxonomic level. Dereplicated high-quality MAGs were used to generate a phylogenetic tree using FastTree (v2.1.11) ^46^ and their relationships drawn with iTOL^47^. CoverM (v0.6.1) ^48^ was used to retrieve MAGs’s relative abundance and read counts.

MAGs were annotated using the NCBI Prokaryotic Genome Annotation Pipeline (PGAP, release 2024-07-18.build7555)^49^ and protein sequences forwarded for functional annotation with EggnNOG-mapper ^50,51^. MAGs were also processed through TORMES ^52^ version 1.3.0 (options --gene_min_id 30 --gene_min_cov 30) to detect antimicrobial resistance (AMR) and virulence genes. For AMR prediction, three databases were adopted by TORMES (ResFinder ^53^, CARD ^54^ and ARG-ANNOT ^55^, whereas virulence genes were detected through the VFDB ^56^. Multi-locus sequence typing (MLST) was performed with the mlst software (Seemann T, mlst Github https://github.com/tseemann/mls) and the PubMLST database ^57^. To investigate molecules potentially associated with microbial and host interactions in the mammary gland, BAGEL4 ^58^ was employed to identify genes predicted to encode bacteriocins in the MAGs, while antiSMASH ^59^, with the strictness parameter set to “relaxed,” was used to predict bioactive compounds as part of their secondary or specialised metabolism.

### 2.6. Statistical analysis

#### 2.6.1. Generalized Additive Model (GAM)

Two GA models were developed to investigate the influence of lactation period (Period, DO-2022, EL-2023, ML-2023, LL-2023), somatic cell count classification (SCC, “High” and “Low”), and individual animal variability (animal) on microbial diversity metrics, specifically Shannon and Simpson indices. Both models were fitted, treating each categorical variable as a random effect by including smooth terms (bs = “re”). The REML method was selected for estimation to optimize the accuracy of parameter estimates and random effect variances. Data handling, model fitting, and visualization were conducted using R (version 4.4.1) with the R packages phyloseq ^60^, mgcv ^61^, and dplyr ^62^.

#### 2.6.2. Metataxonomic analysis

Statistical analysis and graph construction were performed with RStudio (v.1.2.5033) using one or the combination of the following R packages: MicrobiomeR (https://github.com/microbiome/microbiome), dplyr ^62^, ggplot2 (https://ggplot2.tidyverse.org), phyloseq ^60^, tidyr (https://github.com/tidyverse/tidyr), vegan (https://github.com/vegandevs/vegan), and pairwise Adonis (https://github.com/pmartinezarbizu/pairwiseAdonis).

Groups with parametric distribution were analyzed by one-way ANOVA followed by Tukey’s *post hoc*. The Wilcoxon test was used to compare two unpaired groups. Differences were considered significant at P < 0.05. Significant differences in alpha diversity between the groups with high and low SCC, as well as between the different lactation periods, were determined using the alpha function in microbiome R package v.2.1.24, adopting Wilcoxon’s test. For beta diversity, weighted and unweighted UniFrac distances were subjected to permutational multivariate analysis of variance (ANOVA) to assess significant differences (pseudo-F test) in bacterial community composition and structure among the groups with a permutation number of 999. Non-metric multidimensional scaling (NMDS) plots were constructed with microViz ^63^

To investigate the association between microbial taxa and key variables, we employed the MaAsLin 2 ^64^ package with a negative binomial regression model with Benjamini-Hochberg correction (max_significance = 0.25), incorporating both fixed and random effects. The input data matrix included the absolute abundances of microbial taxa at the phylum and genus level, while the metadata comprised variables of interest for the model, including “SCC”, “Period”, and “sampling day”, with “animal” as a random effect.

The resilience of resident microbiota following dysbiosis was evaluated based on a microbial dysbiosis index (MDI), calculated as follows: (MDI_sample_=log(Relative Abundance of non-pathobionts/Relative Abundance of pathobionts). The genera *Staphylococcus*, *Streptococcus*, *Enterococcus*, *Trueperella*, *Klebsiella*, *Serratia*, *Peptostreptococcus*, *Corynebacterium*, and *Escherichia-Shigella* were defined as pathobionts. A quarter was defined as dysbiotic if it exhibited a Shannon diversity index lower than the mean of quarters with somatic cell counts (SCC) below 100,000 cells/mL and an MDI greater than 0, indicating a predominance of dysbiotic microorganisms within the specific sample.

To reduce the dimensionality of the metataxonomic data, t-distributed stochastic neighbor embedding (t-SNE) was run using the R function Rtsne from the Rtsne package ^65^ and selecting the three-dimensionality output option. A hierarchical clustering method using the “hclust” function in R was then used on the output to identify groups of samples with the same microbiota profile.

#### 2.6.3. Microbial differential abundance analyses based on microbial functional profile

To explore associations between microbial features and metadata variables, microbial differential abundance analyses were conducted using the Maaslin2 package ^64^ with negative binomial regression models, incorporating fixed effects (e.g., “Pathogen”, “SCC”, and “Lactation period”), random effects (e.g., “animal”), and Benjamini-Hochberg correction (max_significance = 0.25), without normalization or transformation.

## 3. RESULTS

### 3.1. Changes in SCC and pathogen distribution across lactation periods reveal significant variability

In this study, we collected 342 individual quarter-level milk samples from 24 Norwegian Red dairy cows (8 primiparous and 16 multiparous). These samples were collected during four lactation periods and encompassed two lactation cycles. The pre-drying period in 2022 (DO-2022) was used as the baseline. In 2023, milk samples were collected at three distinct stages of the lactation cycle based on the days in milk (DIM) of each animal. Samples were obtained during early lactation (EL-2023, median DIM: 22 days), mid-lactation (ML-2023, median DIM: 126 days), and late lactation (LL-2023, median DIM: 244 days), corresponding to the respective stages of the lactation cycle.

Based on SCC, a total of 233 samples were classified as low SCC (< 100.000 cells/mL), while 109 samples were categorized as high SCC (> 100.000 cells/mL). The percentage of quarters with low SCC was notably higher during EL-2023 (80%) and ML-2023 (78.9%) compared to DO-2022 (53.1%) and LL-2023 (53%), with high SCC percentages of 20%, 21.1%, 46.9%, and 47%, respectively. As depicted in **Figure 1A**, there was a statistical difference between the different lactation periods, except between DO-2022/LL-2023 and EL-2023/ML-2023. The distribution of individual bacterial counts (IBC) exhibited a comparable pattern, revealing a significant difference between samples classified as low SCC and those categorized as high SCC. Specifically, low SCC samples had a mean IBC of 3.31 ± 0.475, while high SCC samples showed a mean IBC of 3.74 ± 0.334.

**Figure 1.**
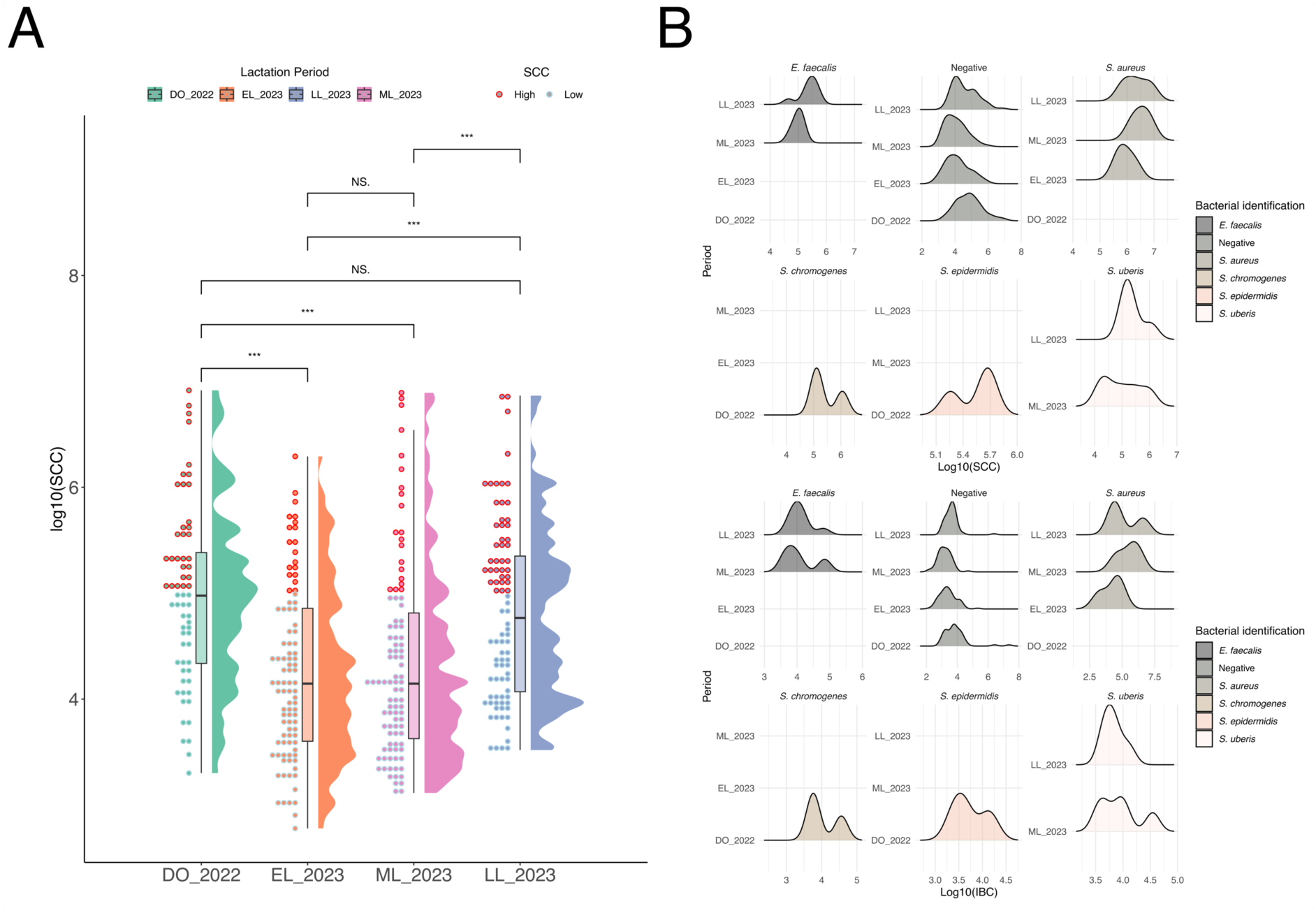
Raincloud plot **(A)** illustrating somatic cell count (log10(SCC)) across different lactation periods (DO-2022, EL-2023, ML-2023, and LL-2023). Individual samples are represented by dots, color-coded as high SCC (red) or low SCC (light blue). Side histograms depict data distribution, while central bars indicate the median and interquartile range. Statistical comparisons between lactation periods are denoted by significance levels: ***p < 0.001, **p < 0.01, and NS (not significant). Density plot **(B)** showing the relationship between SCC and bacterial count (log10(IBC)) across lactation periods, stratified by bacterial species identified using MALDI-ToF. Density curves represent distinct pathogens (*E. faecalis*, *S. aureus*, *S. chromogenes*, *S. epidermidis*, and *S. uberis*) and samples classified as “negative”, highlighting variations in bacterial prevalence and load among lactation periods.

The percentage of quarters testing positive or negative for mastitis pathogens, identified via MALDI-Tof, also reflected the trends observed for SCC and IBC (**Figures 1B and 1C**). The lowest detection of mastitis-causing pathogens was recorded during EL-2023, with only 10.5% of samples testing positive. Among the mastitis-causing pathogens identified in this study (from 60 samples), *E. faecalis* emerged as the predominant etiological agent, accounting for 30% of the cases. *S. aureus*, *S. uberis*, and *S. chromogenes* were detected in 25%, 20%, and 10% of the samples, respectively. The bacterial species *S. epidermidis* (7%), *C. bovis* (3%), *S. dysgalactiae* (3%), and *S. haemolyticus* (2%) were identified to a lesser extent.

The relationship between SCC and IBC ratio concerning specific mastitis-causing pathogens was also investigated. Groups with fewer than three samples, specifically *C. bovis*, *S. dysgalactiae*, and *S. haemolyticus*, were excluded from this analysis. Our findings indicate that the level of somatic cells was significantly higher in samples associated with *S. epidermidis* compared to those classified as negative, with an adjusted p-value of 0.014. This suggests that a small number of *S. epidermidis* cells may be sufficient to trigger a strong pro-inflammatory response in the mammary gland.

When we examined these same samples in relation to bacterial counts on blood agar plates (after 48 hours of incubation at 37°C), alongside individual bacterial count (IBC), somatic cell count (SCC), and the identified pathogen (**Figure 2A**), we observed that many samples classified as negative (i.e.. absence of mastitis-causing pathogens according to the gold standard) still showed high plate counts. Specifically, 129 samples had colony counts ranging between 30 and 300 CFUs. From these samples, 57 showed SCC higher than 100,000 (median: 2.9 x 10^5^ cells/mL) and a median IBC of 7.6 x 10^3^ cells/mL. Our findings indicate that these samples harbor microorganisms distinct from the major mastitis-causing pathogens, which, although not routinely identified, may still contribute to increased somatic cell counts in the mammary gland.

**Figure 2.**
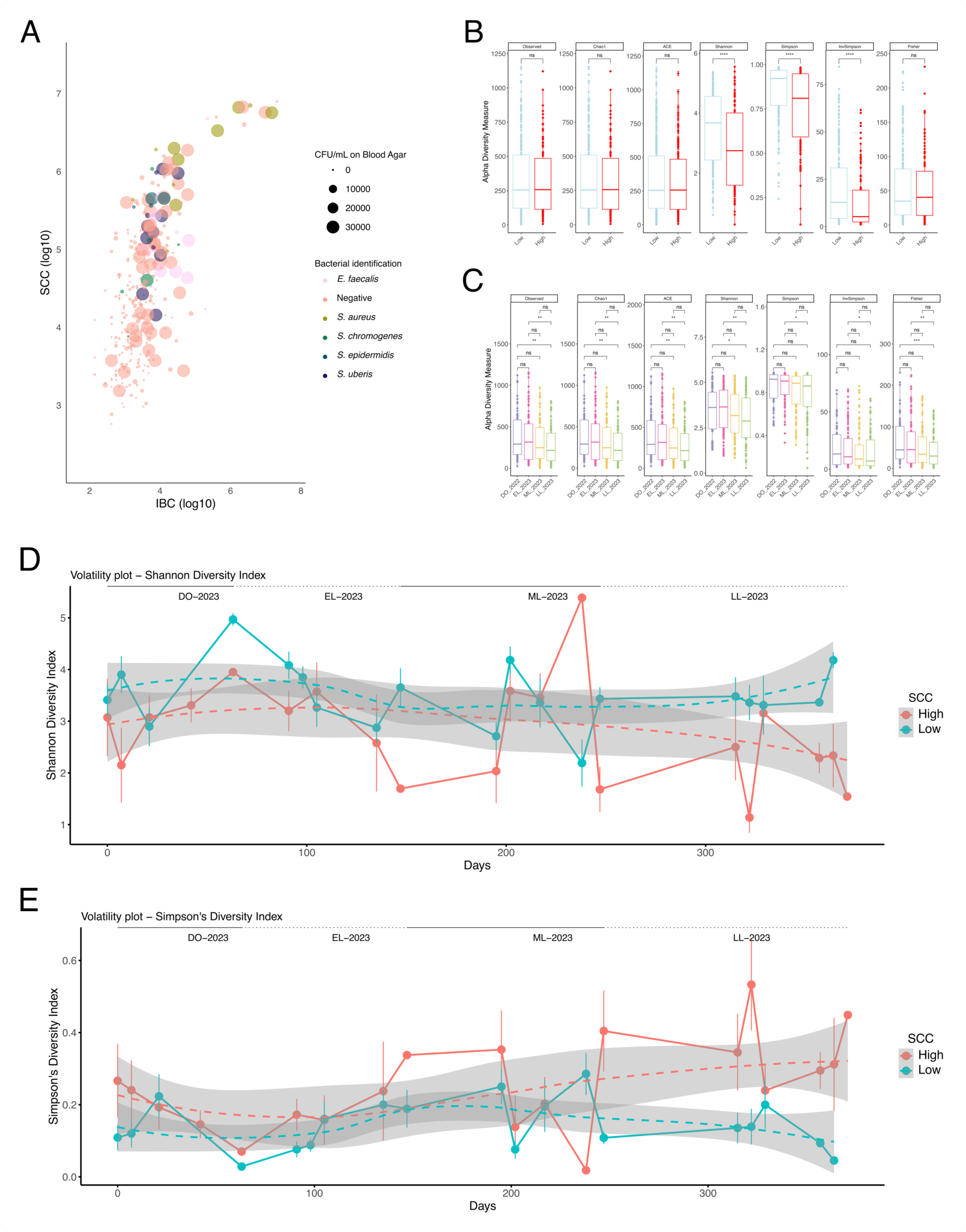
Bubble chart **(A)** displaying the relationship between somatic cell count (SCC) and individual bacterial count (IBC) across samples, categorized by bacterial species. Bubble sizes represent CFU/mL on blood agar, while colors indicate bacterial species (*E. faecalis*, *S. aureus*, *S. chromogenes*, *S. epidermidis*, *S. uberis*), and milk samples classified as “negative” for specific mastitis-causing pathogens. Box plots comparing Shannon **(B)** and Simpson **(C)** diversity indices between high and low SCC groups across different time points and lactation periods. Statistical significance is denoted as ***p < 0.001, **p < 0.01, and *p < 0.05. Volatility plot of Shannon **(D)** and Simpson **(E)** diversity indices over time, showing trends across lactation periods (DO-2023, EL-2023, ML-2023, and LL-2023). Solid and dashed lines represent high and low SCC groups, respectively, with shaded areas indicating 95% confidence intervals with group-wise comparisons highlighted.

### 3.2. Metataxonomic analysis

#### 3.2.1. Alpha and beta diversity analyses show significant effects of lactation period on diversity indices

In this longitudinal study, we investigated the microbial composition of 306 milk samples encompassing a whole lactation cycle. By conducting amplicon sequencing of the hypervariable region V3-V4 of the 16S rRNA gene, we obtained a total of 1.3 × 10^7^ high-quality filtered sequences after the removal of chimeric, contaminants, and spurious sequences (e.g.. mitochondrial and chloroplast sequences). The median number of sequences per sample in the low SCC group was 37,589, while the high SCC group had a median of 47,282 sequences. Across all groups, the overall median number of sequences per sample was 42,970.

We first investigated alpha diversity analysis considering the variables SCC (low and high) and lactation period (DO-2022, EL-2023, ML-2023, and LL-2023). When we analyzed the variable SCC, statistical significance was observed for the indices Shannon, Simpson, and inverse Simpson (**Figure 2B**). Overall, samples with low SCC exhibited greater microbial diversity and lower dominance of individual species. When the different periods were considered, five out of seven indices showed higher diversity measurements for the periods DO-2022 when compared to LL-2023. All seven indices were statistically significant for the pairwise comparison between EL-2023 and LL-2023 (**Figure 2C**). As depicted in the volatility plots (**Figures 2D and 2E**) for the Shannon and Simpson indices, there is a clear trend indicating a reduction in both indices at the end of lactation in samples containing more than 100,000 cells/mL.

We also employed the generalized additive model (GAM) to analyze how the Shannon and Simpson diversity indices were influenced by different predictor variables. The GAM analysis revealed significant effects of the lactation period, somatic cell count, and individual animal variability on both the Shannon and Simpson indices, while bacterial count did not show a significant impact. The smooth term for the lactation period indicated a modest but statistically significant effect on microbial diversity (Shannon: edf = 2.10, F = 2.72, p = 0.039; Simpson: edf = 2.11, F = 2.66, p = 0.047), suggesting variation in diversity across lactation stages. Somatic cell count had a highly significant effect (Shannon: edf = 0.92, F = 28.95, p < 0.001; Simpson: edf = 0.94, F = 38.44, p < 0.001), indicating that the degree of mammary gland inflammation plays a critical role in shaping the microbiota. Additionally, the random effect of individual animals was significant (Shannon: edf = 16.37, F = 2.25, p < 0.001; Simpson: edf = 14.7, F = 1.72, p = 0.002), highlighting considerable inter-animal variability in microbial diversity. However, the bacterial count was not a significant predictor of diversity (Shannon: Estimate = 1.49e-08, p = 0.681; Simpson: Estimate = 4.353e-09, p = 0.387). The model explained 24.9% and 23.4% of the deviance, with adjusted R-squared values of 19.5% and 18.4% for the Shannon and Simpson indices, respectively, underscoring the relevance of the factors considered in the model.

The PERMANOVA analyses using unweighted and weighted UniFrac distances revealed significant effects of various factors on the microbial community structure (**Figure 3**). For the unweighted UniFrac analysis, the lactation period explained 3.01% of the variance in microbiota composition (F = 3.20, p = 0.01), indicating shifts in microbial diversity across different stages of lactation. Individual animal variability also contributed 9.42% to the observed variance (F = 1.25, p = 0.01), highlighting notable inter-animal differences in microbial composition. However, somatic cell count and the history of mastitis-causing pathogens identified in milk samples were not significant predictors, accounting for only 0.30% and 2.85% of the variance, respectively. The majority of the variation, 83.97%, remained unexplained by the measured factors, suggesting other influences on the microbial community.

**Figure 3.**
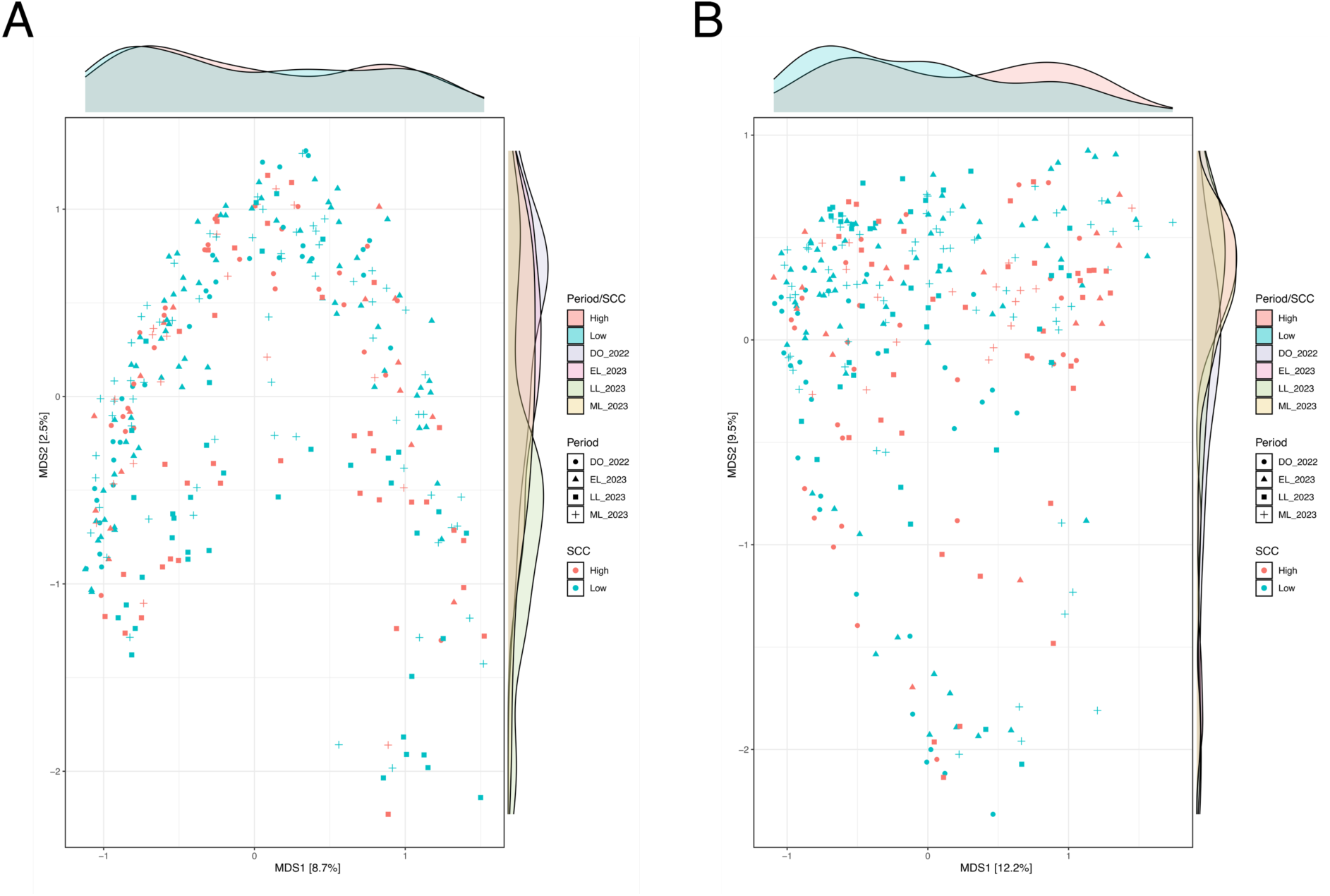
Non-metric multidimensional scaling (NMDS) plot based on unweighted **(A)** and weighted **(B)** UniFrac distances, illustrating microbial community composition across different lactation periods (DO-2022, EL-2023, ML-2023, LL-2023) and SCC levels (high and low). Points represent individual samples, with colors indicating SCC levels and shapes representing lactation periods. Marginal density plots along the axes depict the distribution of samples by MDS dimensions.

In contrast, the weighted UniFrac analysis indicated more pronounced effects of the measured variables. Somatic cell count had a small but significant effect, explaining 0.78% of the variance (F = 3.22, p = 0.01). The lactation period played a more substantial role, explaining 5.52% of the variance (F = 7.57, p = 0.01), reinforcing the influence of lactation stages on microbial composition. Individual animal variability was the largest contributor to variance, explaining 15.73% (F = 2.69, p = 0.01), while the identification of major mastitis-causing pathogens in milk samples contributed 7.17% (F = 3.28, p = 0.01), highlighting the impact of dysbiosis on microbial communities in the udder. The remaining 65.18% of the variance was attributed to residual (i.e., unmeasured factors), reflecting biological complexity and other unknown influences shaping the udder microbiota.

#### 3.2.2. Differential abundance analysis and multivariable associations reveal shifts in *Actinobacteriota* across lactation stages

After clustering at a 99% similarity threshold and preprocessing steps (removal of *g-Mitochondria*, *o-Chloroplast*, and *d-Archaea*), we identified a total of 48,622 ASVs. Four phyla – *Firmicutes* (median: 63%; mean: 60.1%), *Actinobacteria* (median: 20%; mean = 25.2%), *Proteobacteria* (median: 7%; mean: 11.4%), and *Bacteroidota* (1%; mean: 2.3%) – comprised approximately 99% of the milk bacterial population. At the genus level, ten taxa stand out and comprise approximately 65% of the microorganisms at this taxonomic level: *Corynebacterium* (median: 12.9%; mean: 19.4%), *Staphylococcus* (median: 4.0%; mean: 17.0%), *Bradyrhizobium* (median: 1.8%; mean: 5.4%), *Streptococcus* (median: 0.2%; mean: 4.9%), *Aerococcus* (median: 2.6%. mean: 4.8%), *Romboutsia* (median: 3.4%; mean: 4.5%), *Enterococcus* (median: 0.2%; mean: 3.1%), *Kocuria* (median: 0.9%; mean: 2.0%), *Lactococcus* (median: 0.2%; mean: 1.7%), and *Weissella* (median: 0.6%; mean: 1.6%).

We performed a differential abundance analysis using MaAsLin2 to determine multivariable associations across samples grouped by somatic cell count (SCC), lactation period, sampling day, and microbial composition at both the phylum and genus levels. We found significant results for 15 phyla **(Supplementary Table 1)**. Samples with low SCC showed a significantly greater abundance of eight phyla, with *Proteobacteria*, *Actinobacteriota*, and *Bacteroidota* particularly prominent based on their prevalence in our dataset (belonging to the top four phyla). Additionally, shifts in bacterial phyla were observed across different lactation periods. While the abundance of the *Firmicutes* phylum remained relatively stable throughout the lactation periods (q-value > 0.25), we observed an enrichment of the *Proteobacteria* phylum in the early lactation period (EL-2023), which gradually decreased in the subsequent periods. In contrast, members of the *Actinobacteriota* phylum were enriched in the mid- (ML-2023) and late lactation (LL-2023) periods. When considering the continuous variable “day”, we observed a distinct fluctuation in *Actinobacteriota* abundance across the studied periods. Specifically, *Actinobacteriota* was highly abundant during the baseline period DO-2023, but this abundance decreased in the early lactation period (EL-2023), followed by two subsequent increases during the mid-(ML-2023) and late lactation (LL-2023) periods.

Our genus-level analysis identified 66 differentially abundant genera **(Supplementary Table 2)**. Samples with low SCC showed enrichment in several genera, with *Corynebacterium*, *Bradyrhizobium*, *Romboutsia*, and *Lactococcus* standing out among the top 10 genera. Conversely, *Staphylococcus* was significantly enriched in samples with high SCC. Examining differentially abundant genera by lactation period, *Bradyrhizobium*, Unidentified *Lactobacillaceae* family. *Pediococcus*, Unidentified *Aerococcaceae* Family, and *Weissella* were enriched in samples collected during EL-2023 compared to DO-2022. Additionally, two genera from the families *Aerococcaceae* and *Lactobacillaceae* were enriched in ML-2023. When analyzed by sampling day, *Bifidobacterium* and *Bacillus* displayed positive coefficients, indicating a progressive increase in abundance over time, whereas the abundance of Unidentified *Lactobacillaceae* family decreased over time.

### 3.3. The integration between t-SNE and culturomics improves milk microbiota classification

The microbial composition of the 295 samples was subjected to dimensionality reduction t-SNE analysis. An optimal number of k-mean clusters was identified by using the “Elbow method,” and the microbiota was divided into eighteen clusters **(Supplementary Figure 1A-C)**. The eighteen clusters were then used to investigate the microbial composition, which significantly differed in main ASVs and genera. Each cluster was then used as a factor to map Chao1 and Shannon diversity indices and other variables such as SCC and IBC. As depicted in **Figure 4**, cluster analysis revealed that IBC levels varied significantly from the global mean in clusters 9, 10, 13, 14, 16, and 17. For SCC, statistically significant differences were observed in clusters 9, 10, 11, 13, 14, 15, and 17. The diversity indices Chao1 and Shannon also exhibited substantial variation across clusters. For the Chao1 index, clusters 6, 7, 10, 13, 14, 15, 16, and 18 did not differ significantly from the global mean. In contrast, for the Shannon index, all clusters except 2, 7, 9, 10, 15, 16, and 18 showed significant differences from the global mean.

**Figure 4.**
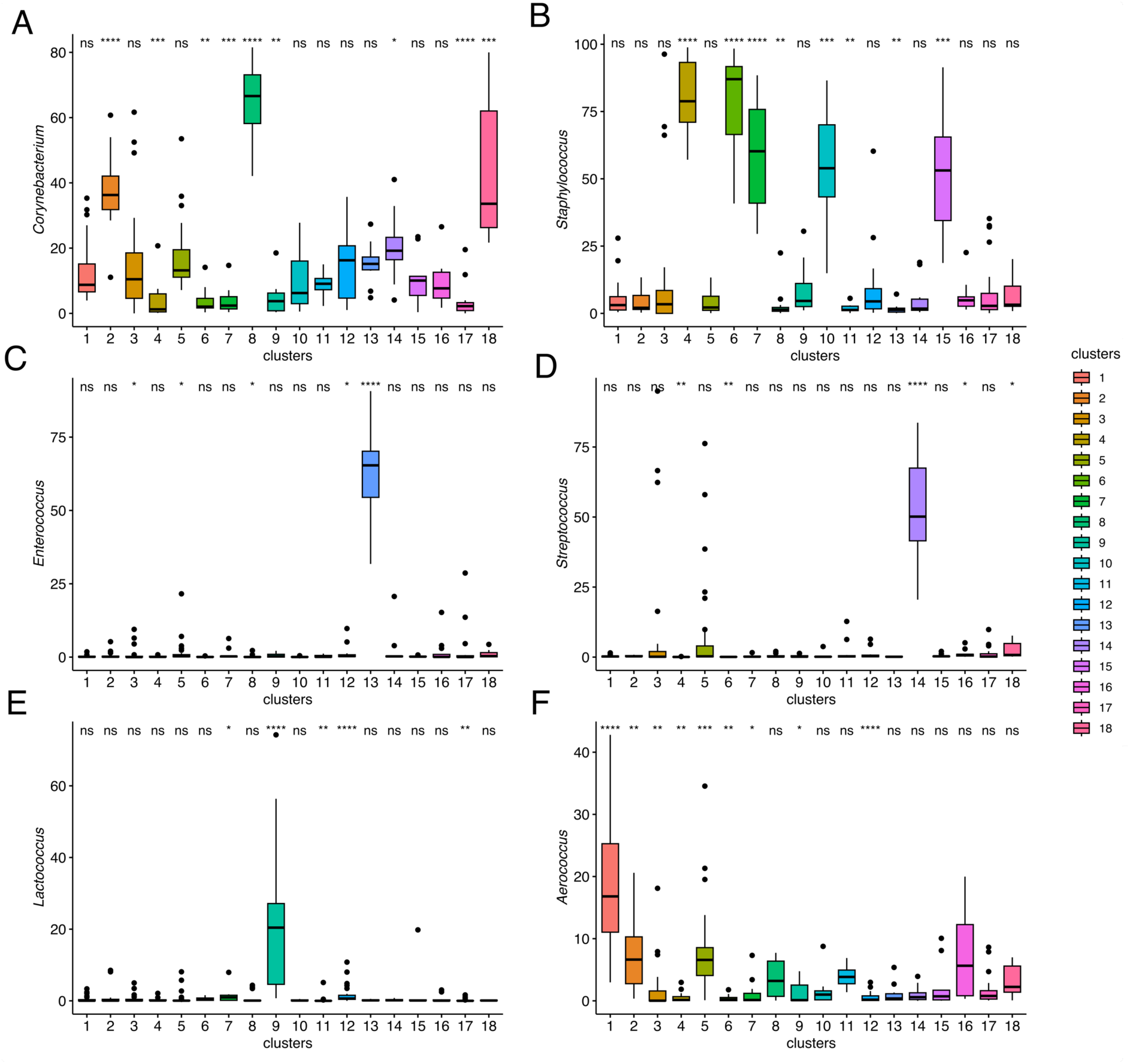
Box plots **(A–F)** showing the distribution of the relative abundance of the six dominant genera (*Corynebacterium*, *Staphylococcus*, *Enterococcus*, *Streptococcus*, *Lactococcus*, and *Aerococcus*) across the 18 microbiota clusters. Each box represents the interquartile range (IQR), with the median indicated by a horizontal line and whiskers extending to 1.5 times the IQR. Outliers are shown as individual points. Statistical significance of differences in relative abundance between clusters and the global mean is denoted above each box plot (p < 0.05; p < 0.01; *p < 0.001; **p < 0.0001; ns = not significant).

We conducted an analysis to examine the distribution of the predominant bacterial genera across different clusters (**Figure 5**). The genera *Corynebacterium*, *Staphylococcus*, and *Aerococcus* show high prevalence and widespread distribution across clusters, indicating that these genera are commonly found across diverse milk samples used in this study and are not restricted to any specific group. In contrast, *Enterococcus* and *Streptococcus* were majority limited to a single cluster each, respectively, clusters 13 and 14.

**Figure 5.**
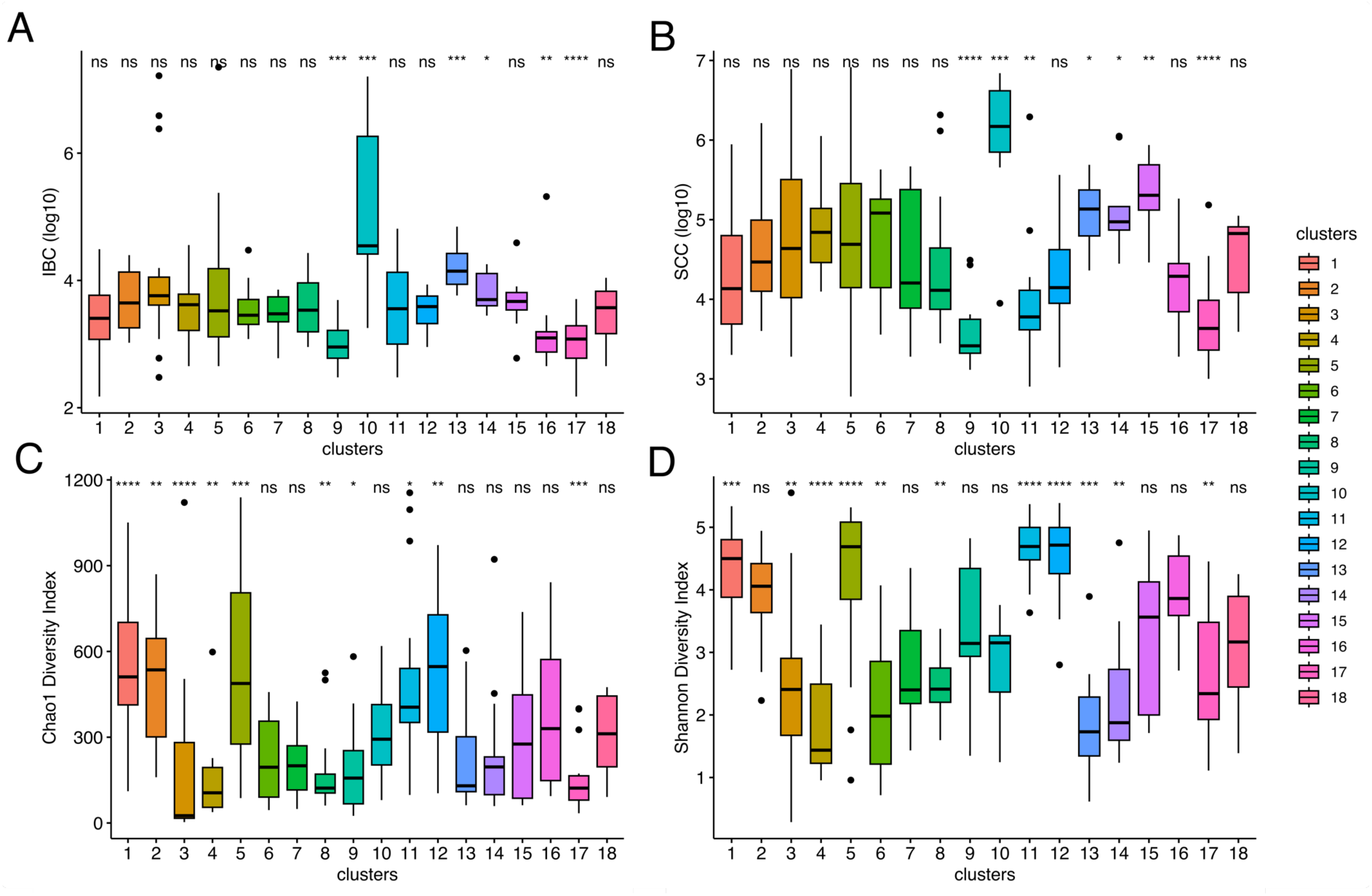
Distribution of bacterial count **(A)**, somatic cell count **(B)**. and alpha diversity indices (**C** and **D**) across the 18 microbiota clusters. Each box represents the interquartile range (IQR), with the median indicated by a horizontal line and whiskers extending to 1.5 times the IQR. Outliers are shown as individual points. Statistical significance of differences in relative abundance between clusters and the global mean is denoted above each box plot (p < 0.05; p < 0.01; *p < 0.001; **p < 0.0001; ns = not significant).

Considering a minimum mean relative abundance of 10% per cluster **(Supplementary Figure 2)**, we observed a notably high relative abundance of *Staphylococcus* in five clusters (cluster 4: 80.50%, cluster 6: 77.88%, cluster 7: 58.85%, cluster 10: 56.96%, cluster 15: 52.69%). Combined with the culturomics’ analysis, cluster 4 was enriched for *S. chromogenes* (4 positive samples out of 12), whereas cluster 10 was enriched for *S. aureus* (7 positive samples out of 9). Only one species of *S. haemolyticus* was identified by MALDI-Tof (cluster 6).

We also conducted a MegaBLAST analysis on the top 20 ASVs to investigate species-level identities within clusters with high prevalence of *Staphylococcus* and *Corynebacterium* and *Streptococcus* (**Figure 6**). For cluster 4, samples showed a homogenous distribution of the four most abundant ASVs, here annotated as *S. chromogenes* (identity scores ranging from 99.53 to 100%). Cluster 6 showed a high abundance of an ASV that had as top hits (identity of 100%) with *S. borealis*, *S.taphylococcus pragensis*, *S.taphylococcus petrasii,* and *S.taphylo croceilyticus*.

**Figure 6.**
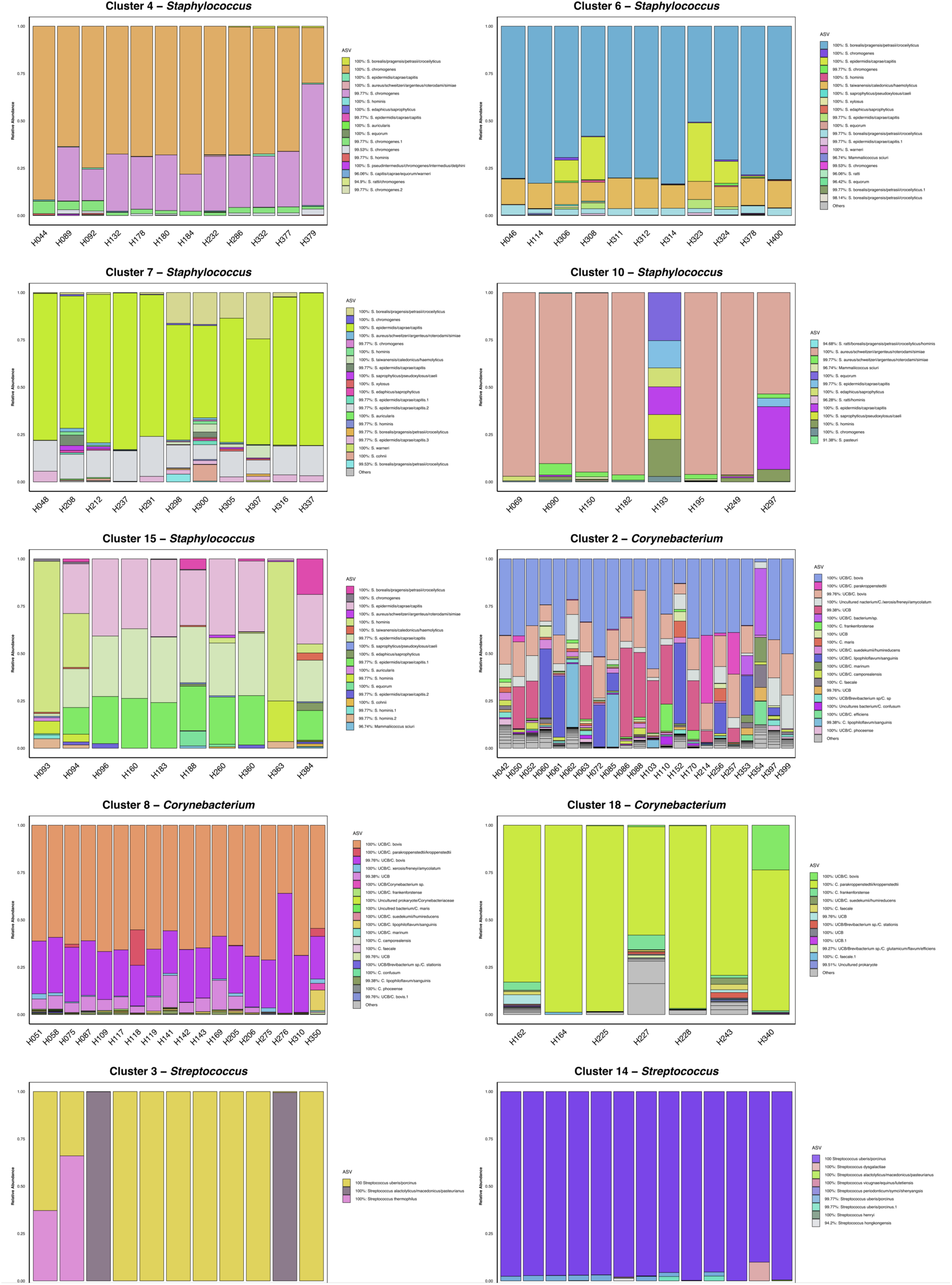
Stacked bar plots illustrating the relative abundance of the most abundant amplicon sequence variants (ASVs) across microbiota clusters. Each bar represents the proportional composition of ASVs within a specific cluster, with colors corresponding to distinct ASVs as shown in the legend.

Cluster 10 was primarily dominated by a single ASV with 100% identity, annotated as *S. aureus*, *S. schweitzeri*, *S. argenteus*, *S. roterodami*, and *S. simiae*. Notably, sample H193 displayed a microbial profile distinct from the other samples in this cluster, diverging from the typical dominance of *S. aureus*. Cluster 7 was dominated by an ASV with 100% identity matching *S. epidermidis*, *S. caprae*, and S. *capitis*. Additionally, four samples (H298, H300, H305, and H307) showed a high abundance of an ASV annotated with 100% identity as *S. borealis*, *S. pragensis*, *S. petrasii*, and *S. croceilyticus*. In the final *Staphylococcus* cluster analyzed (cluster 15), ASVs were annotated as *S. epidermidis, S. caprae, S. capitis,* and *S. hominis*.

Regarding *Corynebacterium*, eleven out of eighteen clusters showed a relative abundance greater than 10%. Clusters 2 (37.68%), 8 (64.92%), and 18 (44.56%) exhibited particularly high abundances. It is important to note that from culturomics’ data, the detection of *Corynebacterium* species was limited to only two samples (H087 and H109), both positive for *Corynebacterium bovis*. The ASV-level investigation provided insights into the composition of these clusters. Our analysis revealed that several ASVs had the best hits with uncultured bacteria. Cluster 2 stood out for its heterogeneity of *Corynebacterium* species, a pattern not observed in clusters 8 and 18, and overall enrichment for *C. bovis*. Cluster 8 showed a more homogeneous distribution of ASVs, and specifically two sequence variants annotated as uncultured bacterium/*C. bovis* (identities of 99.76% and 100%).

Lastly, clusters 3 and 14 showed a high abundance of members of the genus *Streptococcus*. Cluster 3 was enriched for three ASVs (100% identity) annotated as *Streptococcus uberis* /*Streptococcus porcinus*, *Streptococcus alactolyticus*/*Streptococcus macedonicus*/*Streptococcus pasteurianus*, whereas cluster 14 was dominated by a single ASV identified as *Streptococcus uberis* (identity: 100%).

#### 3.3.1. Udder microbiota dynamics reveal persistent infection by *S. aureus* and *S. chromogenes*, but not *S. uberis*

Following t-SNE analysis to enhance sample classification and elucidate underlying patterns in the dataset, we conducted a quarter-level analysis to track temporal fluctuations in genus-level taxa before and after detecting the pathogen identified by the gold standard method. We focused on samples testing positive for Staphylococci, Enterococci, and Streptococci. Notably, these milk samples predominantly grouped into clusters 4, 10, 13, and 14. For robustness, only quarters with a minimum of three sampling points were included in the analysis (**Figure 7**). To estimate the resilience of resident microbiota following a dysbiosis event, we established a microbial dysbiosis index combined with the Shannon diversity index. A quarter was classified as dysbiotic if it exhibited a Shannon index below 3.5 (the mean value for quarters with somatic cell counts below 100,000 cells/mL) and an MDI greater than 0, indicating a predominance of dysbiotic microorganisms within that sample.

**Figure 7.**
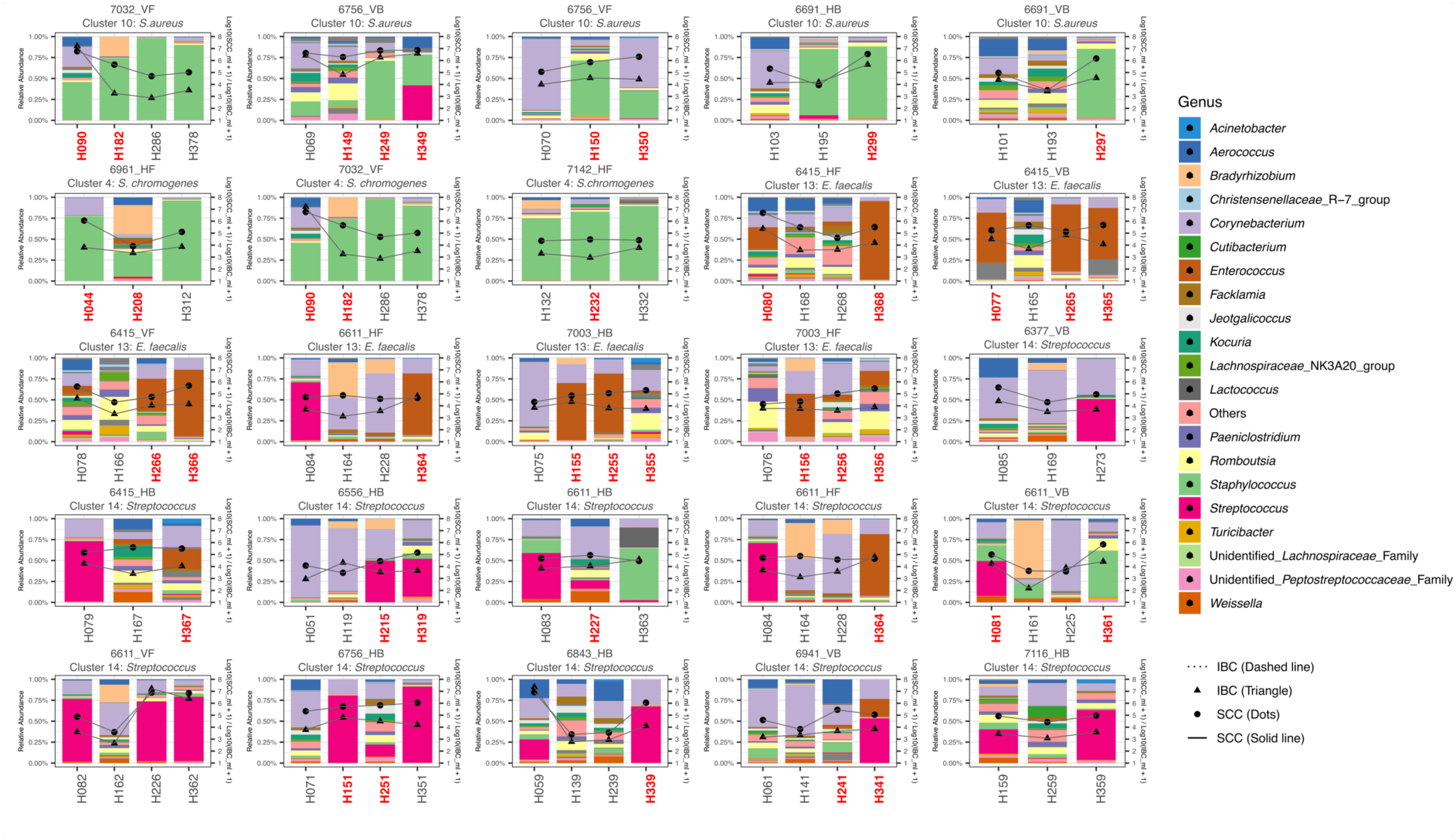
Udder microbiota dynamics. Stacked bar plots depicting the relative abundance of the 20 most abundant genera across different sampling quarters. Each bar represents the proportional composition of genera, with colors corresponding to specific genera as indicated in the legend. Genera with lower abundances are grouped under “Others.” Bacterial and somatic cell counts are overlaid: triangles connected by dashed lines represent IBC levels, while dots connected by solid lines indicate SCC levels, illustrating changes in microbiota composition in relation to these metrics over sampling time. The analyzed animal and sampling quarter, along with microbiota clusters, are labeled above each panel, with sample identification numbers displayed below each plot. Samples highlighted in red indicate the detection of a mastitis-causing pathogen identified via MALDI-ToF analysis.

For cluster 10, which was identified as enriched with *S. aureus*, 5 quarters were analyzed and the longitudinal analysis revealed a persistent state of infection by *S. aureus*. Once an infection was established, *S. aureus* remained consistently present in samples regardless of the lactation stage. Although Intramammary bacterial counts varied over time, somatic cell counts consistently remained elevated, typically exceeding, or in a few cases, hovering around 100,000 cells/mL. A marked depletion of specific taxa, such as *Corynebacterium*, *Aerococcus*, and *Romboutsia*, was observed, along with reductions in other low-abundance taxa grouped here as “Others.” Additionally, a co-infection with species of the genus *Streptococcus* was detected at two specific time points (samples H195 and H349). When the MDI index was calculated for samples in this cluster, it was evident that samples following an episode of high *S. aureus* abundance did not return to a state of eubiosis.

Cluster 4 showed enrichment for *S. chromogenes* and persistent infection behavior was also observed. However, it’s noteworthy that the infection-associated somatic cell counts and intramammary bacterial counts in this cluster are significantly lower than those found in cluster 10. In the case of quarter 7032_VF, both *S. aureus* and *S. chromogenes* were isolated and identified. Interestingly, once this quarter was infected by both species, the microbiota failed to revert to its original state. This phenomenon is also clearly demonstrated in quarters 6691_HB and 6691_VB, which suggests that *S. aureus* and *S. chromogenes* exhibit specific competitive mechanisms that are reflected in a pattern of microbiota depletion. This pattern impedes the recolonization of the native microbiota, leading to a more prolonged dysbiosis status, as evidenced by the dysbiosis indices calculated for those samples.

The analysis of cluster 13, enriched for *E. faecalis*, revealed a comparatively less severe dysbiosis than that observed in quarters infected with *S. aureus* and *S. chromogenes*. In this cluster, a reduction in the relative abundance of taxa such as *Corynebacterium*, *Aerococcus*, and *Romboutsia* was noted. However, the native microbiota appears capable of re-establishing itself, a resilience not observed in clusters 4 and 10. This phenomenon could be observed for the quarters 6415_VB (sample H077: Shannon = 1.73, MDI = 1.03; sample H165: Shannon = 4.77, MDI = - 2.35), 7003_HB (sample H255: Shannon = 2.53, MDI = 1.23; sample H355: Shannon = 5.17, MDI = -0.72), and 7003_HF (sample H156: Shannon = 2.04, MDI = 1.05; sample H256: Shannon = 4.43, MDI = -0.49).

For cluster 14 (characterized by enrichment in *Streptococcus uberis*), we observed that the reduction in *Corynebacterium* was not as pronounced as in clusters dominated by *Staphylococcus* and *Enterococcus*. Although *S. uberis* induced a dysbiotic state in the mammary gland, we identified five instances in which a dysbiotic quarter returned to a eubiotic state, and one which we classified as in the borderline of a eubiotic state (samples H084 and H164), including quarters such as 6611_HF (sample H082: Shannon = 1.65, MDI = 2.33; sample H162: Shannon = 3.92, MDI = - 0.65), 6415_HB (sample H079: Shannon = 1.60, MDI = 2.04; H167 = Shannon = 4.58, MDI = -1.94), 6611_HB (sample H083: Shannon, 2.44, MDI: 1.98), 6611_HF (sample H084: Shannon = 2.36, MDI = 1.55; sample H164: Shannon = 2.96, MDI = -0.21), and 6941_VB (sample H141: Shannon = 3.38, MDI = 0.34; sample H241: Shannon = 4.10, MDI = -0.65).

### 3.4. Metagenomic shotgun sequencing uncovers pathogen-specific metabolic signatures in the bovine mammary gland

After trimming, quality filtering, and in silico separation of bacterial reads from bovine sequences, we obtained approximately 269.5 GB of high-quality PE reads across 73 samples, which were then processed for functional profiling using HUMAnN3. This profiling identified 289 MetaCyc metabolic pathways, which were then assessed for differential abundance with MaAsLin2 considering “SCC” and “pathogen” as fixed effects. At first glance, the heatmap (**Figure 8A**) constructed with the 289 predicted MetaCyc pathways revealed that the samples did not form well-defined clusters when grouped by SCC or lactation period, highlighting a significant degree of heterogeneity. However, four distinct clusters of pathogen-negative samples are evident in the heatmap, characterized by a notable lack of metabolic pathway enrichment. Additionally, clusters of samples identified as positive for *S. aureus* and *E. faecalis* were also discernible.

**Figure 8.**
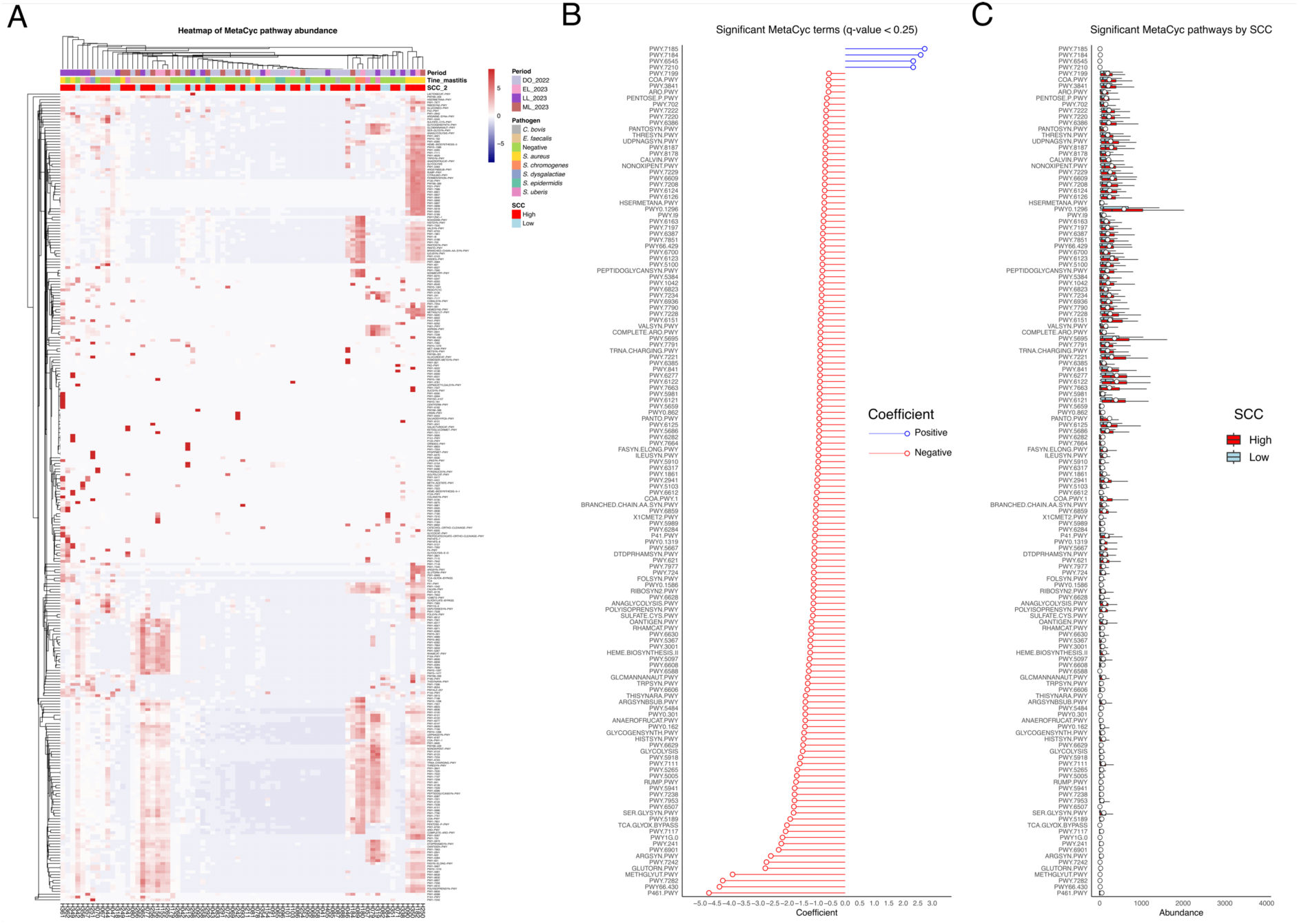
Heatmap **(A)** showing the most abundant MetaCyc pathways detected across 73 metagenomic samples analyzed using HUMAnN3. Hierarchical clustering was applied to both samples and pathways, with samples annotated by lactation period, somatic cell count, and pathogen (identified by MALDI-Tof). Pathway abundance is represented by a red-to-blue color gradient, where darker red indicates higher abundance and darker blue indicates lower abundance. Bar plots **(B)** display the significant MetaCyc pathways (q-value < 0.25) associated with high and low SCC groups, with coefficients categorized as positive (blue) or negative (red) to illustrate their direction of association. Boxplots **(C)** depict the relative abundance of significant MetaCyc pathways stratified by SCC levels, highlighting pathway abundance differences between the two groups.

Focusing on pathways that were differentially expressed (q-value < 0.25) based on somatic cell content, we observed an enrichment of 134 microbial MetaCyc pathways for samples with high SCC and four MetaCyc pathways for samples with low SCC (**Figure 8BC**). When pathways were stratified by bacterial species within the high SCC group, many pathways were common across several bacterial species, including Staphylococci and Streptococci, and were largely associated with central metabolism. Notably, the pathway PWY-6507 (PWY-6507: 4-deoxy-L-threo-hex-4-enopyranuronate degradation) was significantly enriched in *E. faecalis*. In contrast, the four pathways enriched in the low SCC group (PWY-7185, PWY-7184, PWY-6545, and PWY-7210) were primarily related to pyrimidine biosynthesis.

We also assessed the enrichment of metabolic pathways by bacterial species identified via MALDI-ToF, compared to samples classified as negative. This analysis aimed to identify potential metabolic distinctions associated with pathogen presence and uncover infection-specific metabolic signatures unique to the mammary gland. In total, five pathogens were analyzed: *S. aureus*, *S. chromogenes*, *E. faecalis*, *S. epidermidis*, and *S. uberis* (**Figure 9**).

**Figure 9.**
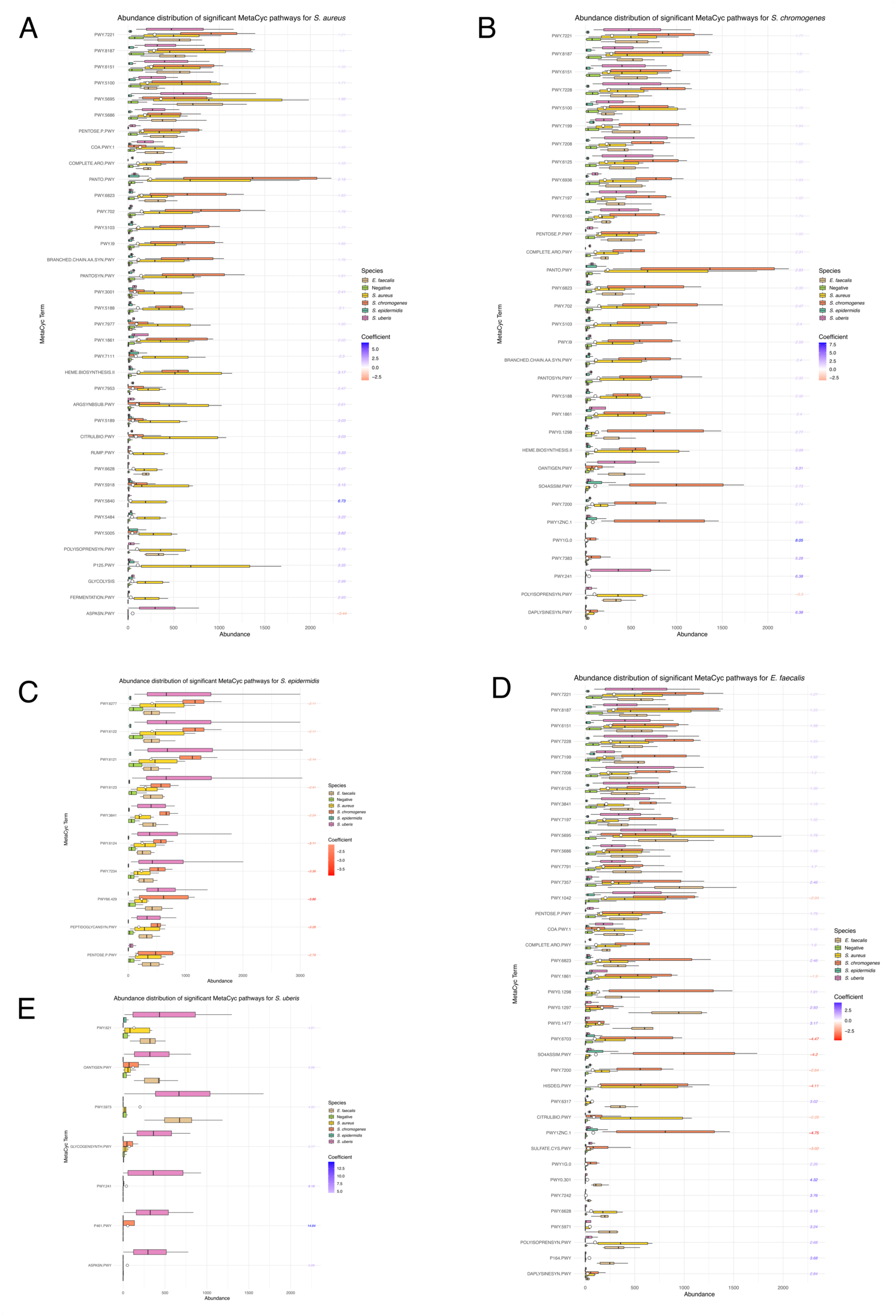
Abundance distribution of significant MetaCyc pathways stratified by bacterial species: (A) *S. aureus*, (B) *S. chromogenes*, (C) *S. epidermidis*, (D) *E. faecalis*, and (E) *S. uberis*. Pathway abundances are shown as boxplots, grouped by species, with pathways color-coded according to their coefficients. Positive and negative coefficients are represented by gradients of blue and red, respectively, highlighting pathways significantly associated with each bacterial species. An empty white dot indicates the mean abundance for each pathway.

For *S. aureus*, 37 pathways were significantly identified, with 36 pathways enriched in this group and one pathway (ASPASN-PWY) more abundant in the reference group. For *S. chromogenes*, 33 pathways were detected, including 32 enriched in this group and one (POLYISOPRENSYN.PWY) enriched in the reference group. In *E. faecalis*, 38 pathways were identified, of which 29 were enriched in this group, while 9 were highly represented in the negative group: PWY-1861, PWY-1042, CITRULBIO-PWY, PWY-7200, SULFATE.CYS.PWY, HISDEG.PWY, SO4ASSIM.PWY, PWY-6703, and PWY1ZNC.1. For *S. epidermidis*, only pathways enriched in the negative group were found, including PWY66-429, PEPTIDOGLYCANSYN.PWY, PWY-6124, PWY-7234, PENTOSE.P.PWY, PWY-3841, PWY-6123, PWY-6122, PWY-6277, and PWY-6121. Lastly, for *S. uberis*, all seven identified pathways were enriched in this group. Overall, these findings highlight distinct metabolic profiles associated with specific pathogens, underscoring the utility of pathway-based analyses to better understand pathogen-host interactions and their metabolic underpinnings in mastitis.

Based on the previously identified pathways with a q-value < 0.25, a Venn diagram was constructed to identify unique and shared MetaCyc pathways (**Figure 10**). For *S. aureus*, 14 unique MetaCyc pathways were identified (PWY-5484, PWY-5005, ARGSYNBSUB.PWY, GLYCOLYSIS, PWY-5840, PWY-7111, PWY-7953, PWY-3001, PWY-7977, FERMENTATION.PWY, PWY-5189, P125-PWY, PWY-5918, and RUMP.PWY). The enrichment of these pathways demonstrated a versatile metabolic profile, encompassing central carbon metabolism, amino acid and nucleotide biosynthesis, stress tolerance, and virulence factors, highlighting the adaptability of *S. aureus* and the potential for pathogenicity in various host environments. *S. chromogenes* exhibited unique metabolic pathways associated with amino acid biosynthesis (PWY-7383, glutamate and glutamine biosynthesis), lipid metabolism (PWY-6936, fatty acid biosynthesis), and energy production (PWY-6163, fermentation), reflecting its ability to adapt to resource-limited environments and its potential role in infection dynamics. The *S. uberis* group showed enrichment in pathways related to carbohydrate metabolism, including glycogen biosynthesis (GLYCOGENSYNTH.PWY), lactose and galactose utilization (P461-PWY), and nucleotide sugar metabolism (PWY-5973), as well as processes involved in cell wall biosynthesis (PWY-621), indicating its capacity to thrive in nutrient-rich environments, such as the mammary gland during infection. For *S. epidermidis,* eight unique metabolic pathways were annotated for this species and involved, for instance, in energy production (glycolysis – PWY66-429), cell-wall structural maintenance (peptidoglycan biosynthesis – PEPTIDOGLYCANSYN-PWY), stress tolerance (polyamine biosynthesis – PWY-6122), and nutrient acquisition (siderophore biosynthesis – PWY-7234; and arginine pathways – PWY-6123). These features contribute to its resilience and success as a commensal and opportunistic pathogen. Finally, *E. faecalis* displayed a metabolic profile enriched in pathways related to amino acid biosynthesis (PWY-5971, PWY.7357), nucleotide metabolism (PWY.6317, P164.PWY), and lipid-related processes (PWY0.1297, PWY0.301), as well as pathways supporting secondary metabolite production (PWY0.1477) and cofactor biosynthesis (PWY.7242, PWY.7791), underscoring its metabolic flexibility and potential resilience in diverse environmental conditions.

**Figure 10.**
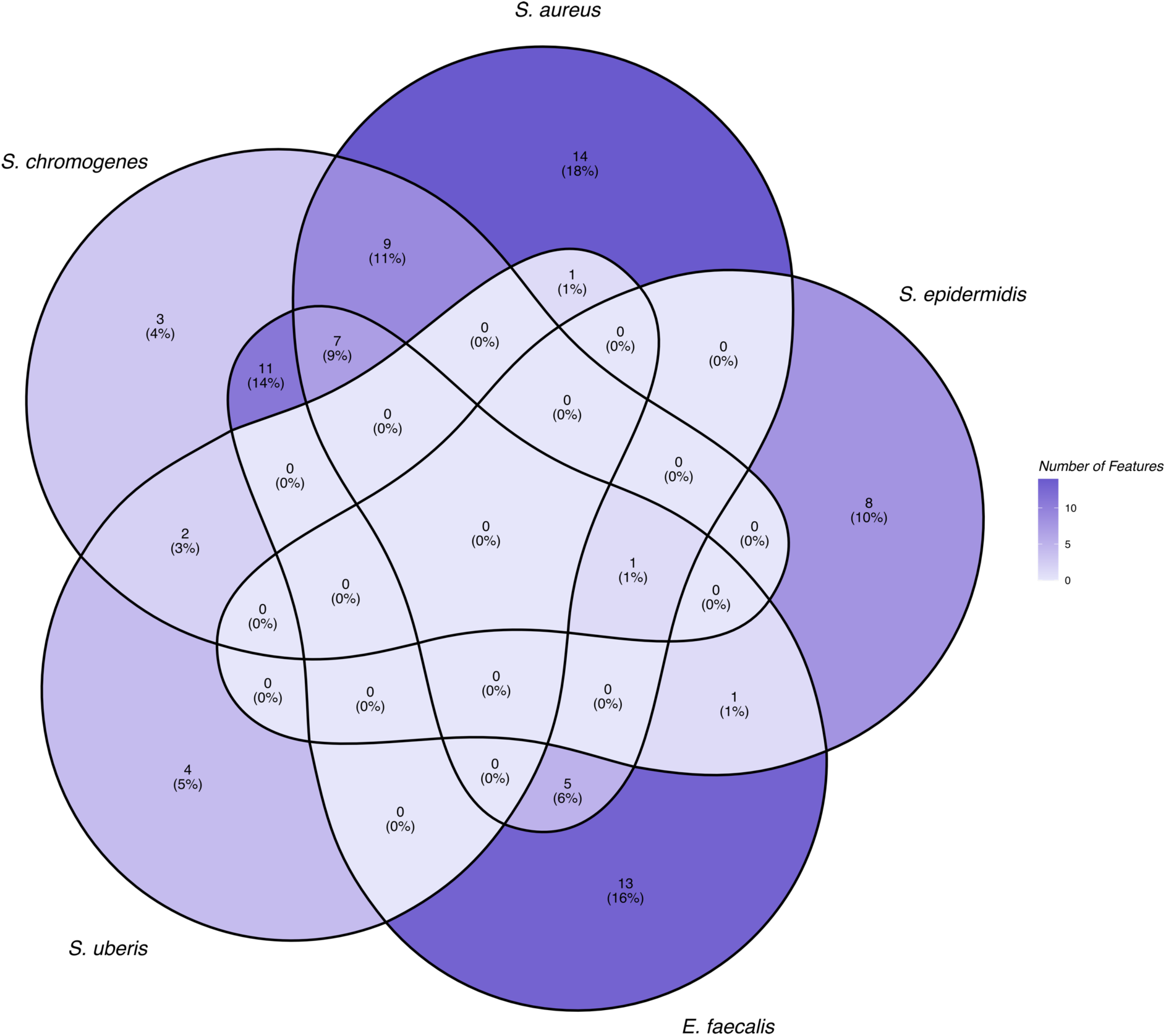
Shared and unique MetaCyc pathways across microorganisms. The Venn diagram shows the number and percentage of pathways that are unique or shared among different species. The shading gradient reflects pathway abundance, with darker shades indicating higher values.

### 3.5. Genome-resolved metagenomic data analysis

#### 3.5.1. MAG quality assessment

The genome-centric approach applied to shotgun reads from milk microbiomes facilitated the reconstruction of 142 metagenome-assembled genomes (MAGs), of which 26 were obtained using a co-assembly method and 116 using individual assembly method. Among these, 38 genomes were classified as high-quality (completion > 90%, contamination < 5%), 50 as medium-quality (completion ≥ 50%, contamination < 10%), and 39 as low-quality (completion < 50%, contamination > 10%) **(Supplementary Table 3)**. MAGs with contamination higher than 10% were excluded from subsequent analyses.

Taxonomic assignments were consolidated at the species level when always possible. For high-quality MAGS, species were annotated as *Corynebacterium bovis* (n = 1), *Corynebacterium kroppenstedtii* (n= 6), *Enterococcus faecalis* (n =9), *Staphylococcus aureus* (n = 5), *Staphylococcus chromogenes* (n = 4), *Staphylococcus epidermidis* (n = 2), *Staphylococcus haemolyticus* (n = 2), *Streptococcus equinus* (n = 2), and *Streptococcus uberis* (n = 7). For medium-quality MAGs, *Corynebacterium bovis* (n = 5), *Corynebacterium kroppenstedtii* (n = 12), *Corynebacterium urogenitale* (n = 1), *Dermatophilus* spp. (n = 2), *Enterococcus faecalis* (n = 10), *Leuconostoc mesenteroides* (n = 1), *Staphylococcus aureus* (n = 2), *Staphylococcus borealis* (n = 2), *Staphylococcus chromogenes* (n = 3), *Staphylococcus epidermidis* (n = 3), *Staphylococcus hominis* (n = 2), *Streptococcus bovis* (n = 1), *Streptococcus dysgalactiae* (n = 2), *Streptococcus equinus* (n = 1), *Streptococcus gallolyticus* (n = 1), and *Streptococcus uberis* (n = 2).

The highest diversity of MAGs were found among the low-quality ones, such as *Aerococcaceae* (n = 2), *Cellulomonas shaoxiangyii* (n = 1), *Corynebacterium bovis* (n = 2), *Corynebacterium glutamicum* (n = 2), *Corynebacterium kroppenstedtii* (n = 4), *Corynebacterium rouxii* (n = 1), *Corynebacterium urogenitale* (n = 1), *Corynebacterium vitaeruminis* (n = 1), *Enterococcus faecalis* (n = 2), *Enterococcus faecium* (n = 1), *Facklamia miroungae* (n = 1), *Kocuria atrinae* (n = 2), *Kocuria carniphila* (n = 1), *Lysobacter* (n = 1), *Micrococcaceae* bacterium (n = 1), *Peptoniphilus* (n = 1), *Peptoniphilus nemausensis* (n = 1), *Ruminococcus flavefaciens* (n = 1), *Ruoffia tabacinasalis* (n = 1), *Staphylococcus aureus* (n = 1), *Staphylococcus chromogenes* (n = 4), *Staphylococcus haemolyticus* (n = 1), *Staphylococcus hominis* (n = 1), *Streptococcaceae* (n = 1), *Streptococcus equinus* (n = 1), *Streptococcus uberis* (n = 1), and *Trichococcus flocculiformis* (n = 1).

After genome dereplication and the exclusion of MAGs with contamination levels exceeding 10% (four MAGs assigned to the family *Aerococcaceae* and one to the family *Micrococcaceae*), 26 MAGs were selected as representative of the metagenomes. These MAGs were subsequently used to map the reads and estimate their relative abundances across the samples (**Figure 11**). The MAG assigned as *Cutibacterium acnes* was identified as a putative contaminant in our dataset and removed from downstream analysis.

**Figure 11.**
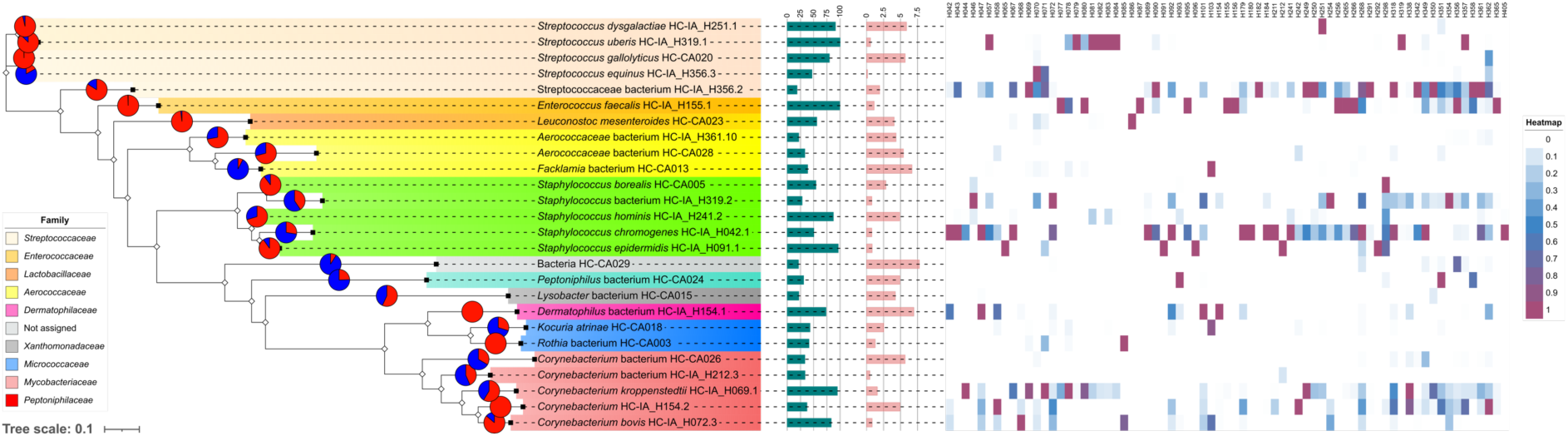
Phylogenetic tree and relative abundance of the species. From left to right tree shows pie charts with species’s abundance in samples with Low (blue) and High (red) somatic cell count, phylogenetic relationships colored at the family level, barplots reporting genome completeness/contamination and heatmap representing relative abundance of species per sample.

To explore the functional potential of the microbiome in greater detail, we predicted microbial genes from the MAGs of interest using the NCBI Prokaryotic Genome Annotation Pipeline, enabling robust downstream comparisons. The resulting protein sequences were annotated using EggNOG, and the COG categories were normalized to the total number of proteins per MAG. Our analysis focused on the bacterial families *Streptococcaceae*, *Aerococcaceae*, *Staphylococcaceae*, and *Corynebacteriaceae* due to their significant representation within the dataset.

As shown in **Figure 12**, the top 20 COG categories primarily comprised proteins associated with metabolism (E: Amino acid transport and metabolism, P: Inorganic ion transport and metabolism, G: Carbohydrate transport and metabolism, C: Energy production and conversion, F: Nucleotide transport and metabolism, H: Coenzyme transport and metabolism, I: Lipid transport and metabolism, and Q: Secondary metabolites biosynthesis, transport, and catabolism), cellular processes and signaling (M: Cell wall/membrane/envelope biogenesis, V: Defense mechanisms, O: Post-translational modification, protein turnover, and chaperones, D: Cell cycle control, cell division, chromosome partitioning, T: Signal transduction mechanisms, and U: Intracellular trafficking, secretion, and vesicular transport), and information storage and processing (L: Replication, recombination, and repair, J: Translation, ribosomal structure, and biogenesis, and K: Transcription). Notably, a substantial proportion of proteins across all bacterial families fell into the “function unknown” group (COG category S), highlighting the large fraction of poorly characterized proteins within these genomes.

**Figure 12.**
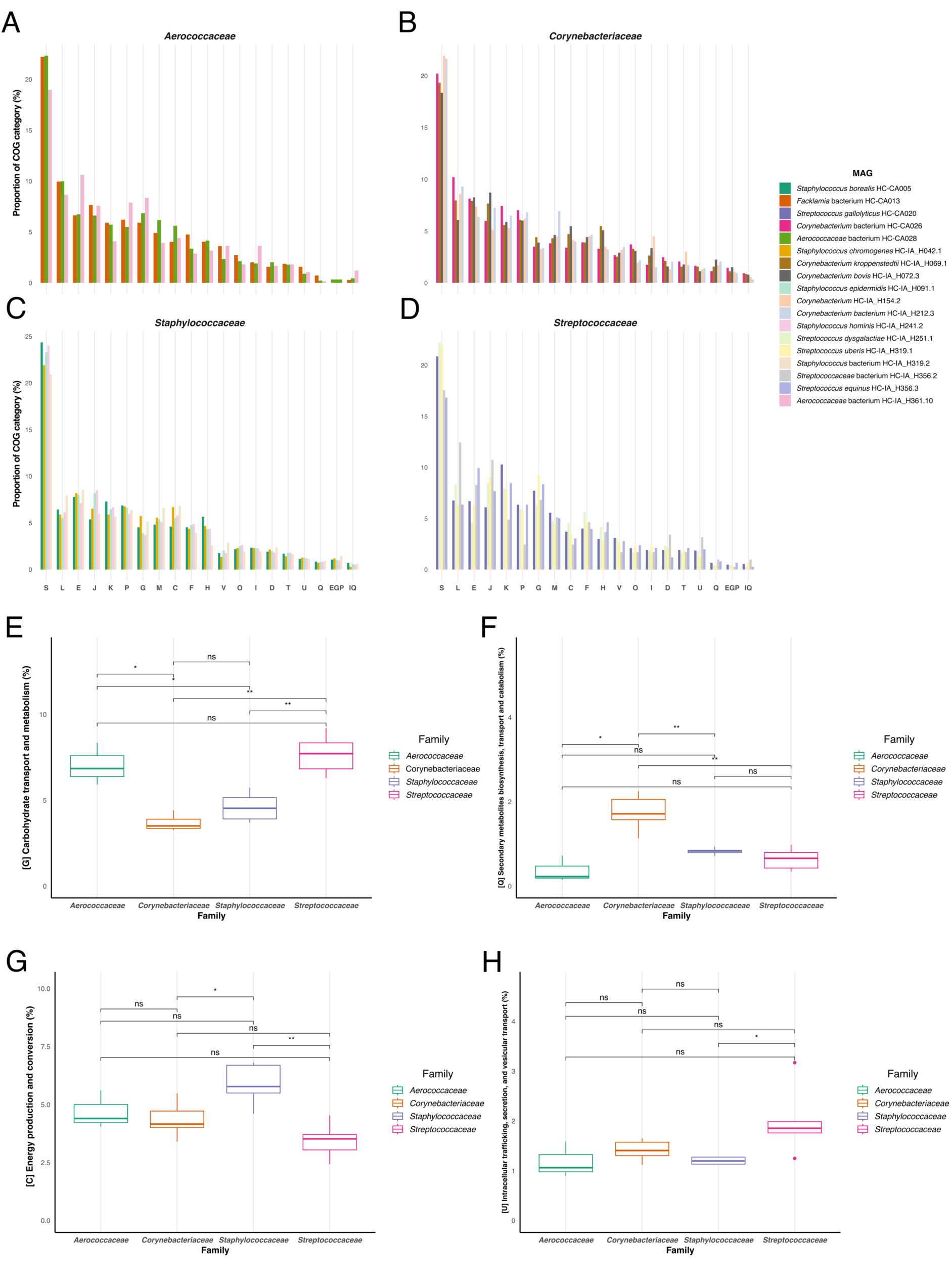
Comparative analysis of clusters of orthologous groups categories among different bacterial families. Bar plots represent the top 20 COG categories across all samples for the bacterial families: *Aeroococcaceae* **(A)**, *Corynebacteriaceae* **(B)**, *Staphylococcaceae* **(C)**, and *Streptococcaceae* **(D)**. Each color indicates a specific metagenome-assembled genome as shown in the legend. COG categories - (S: Function unknown, L: Replication, E: Amino acid transport and metabolism, J: Translation, ribosomal structure and biogenesis, K: Transcription, P: Inorganic ion transport and metabolism, G: Carbohydrate transport and metabolism, M: Cell wall/membrane/envelope biogenesis, C: Energy production and conversion, F: Nucleotide transport and metabolism, H: Coenzyme transport and metabolism, V: Defense mechanisms, O: Posttranslational modification, protein turnover, chaperones, I: Lipid transport and metabolism, D: Cell cycle control, cell division, chromosome partitioning, T: Signal transduction mechanisms, U: Intracellular trafficking, secretion, and vesicular transport, Q: Secondary metabolites biosynthesis, transport and catabolism – are represented on the x-axis, and their normalized proportion on the y-axis. Boxplots **(E–H)** represent COG categories that differed significantly among bacterial families based on the Kruskal-Wallis test. Significant pairwise differences are indicated with * (p<0.05) and ns (not significant).

We aimed to determine which functional categories displayed statistically significant differences in the abundance of annotated proteins across the bacterial families. In total, four COG categories exhibited significant variation. For COG category G (Carbohydrate transport and metabolism), the families *Aerococcaceae* and *Streptococcaceae* demonstrated a higher abundance of orthologs compared to *Corynebacteriaceae* and *Staphylococcaceae*. In COG category Q (Secondary metabolites biosynthesis, transport, and catabolism), *Corynebacteriaceae* displayed the highest abundance of orthologs among the four families. For COG category C (Energy production and conversion), the family *Staphylococcaceae* was enriched compared to *Corynebacteriaceae* and *Streptococcaceae*, although no significant difference was observed when compared to *Aerococcaceae*. Finally, for COG category U (Intracellular trafficking, secretion, and vesicular transport), the family *Streptococcaceae* exhibited a greater abundance of orthologs compared to *Staphylococcaceae*. Overall, the classification of COGs into functional categories provided valuable insights into the metabolic preferences of these bacterial families, highlighting differences in their functional potential and adaptation to specific ecological niches.

#### 3.5.2. Multi-Locus Sequence Typing (MLST), Virulence Factors (VFs) and Antimicrobial Resistance Genes (AMRGs)

All metagenomes, along with high- and medium-quality MAGs, were subjected to MLST analysis. Results revealed that four bacterial species were associated with a single sequence type (*S. aureus*: ST133; *E. faecalis*: ST40; *S. dysgalactiae*: ST302; *S. uberis*: ST1409). In contrast, two sequence types were identified for *S. epidermidis* (ST100 and ST575), while *S. chromogenes* displayed greater diversity with three sequence types (ST1, ST59, and ST104).

The distribution of virulence factors (VFs) across multiple bacterial taxa found in milk samples was analyzed to assess their biosynthetic and pathogenic potential (**Figure 13**). With regards to *Corynebacterium* spp., *C. bovis* exhibited a predominant focus on regulatory factors, representing 51.85% of its VF genes, followed by stress survival and other factors (18.52% each), and nutritional/metabolic factors at 11.11%. Similarly, *C. kroppenstedtii* showed a strong emphasis on regulatory factors (73.33%), with other VF categories accounting for the remaining 26.67%. In *C. urogenitale*, regulation (66.67%) and other factors (33.33%) were the only VF categories identified. Bacterial species annotated as *E. faecalis* showed a broad distribution of VF categories, with adherence genes comprising the largest fraction (43.90%). Exoenzymes (15.70%), immune modulation (12.50%), and stress survival (12.50%) were also well-represented. Other categories included biofilm formation (11.34%), nutritional/metabolic factors (3.78%), and effector delivery systems (0.29%).

**Figure 13.**
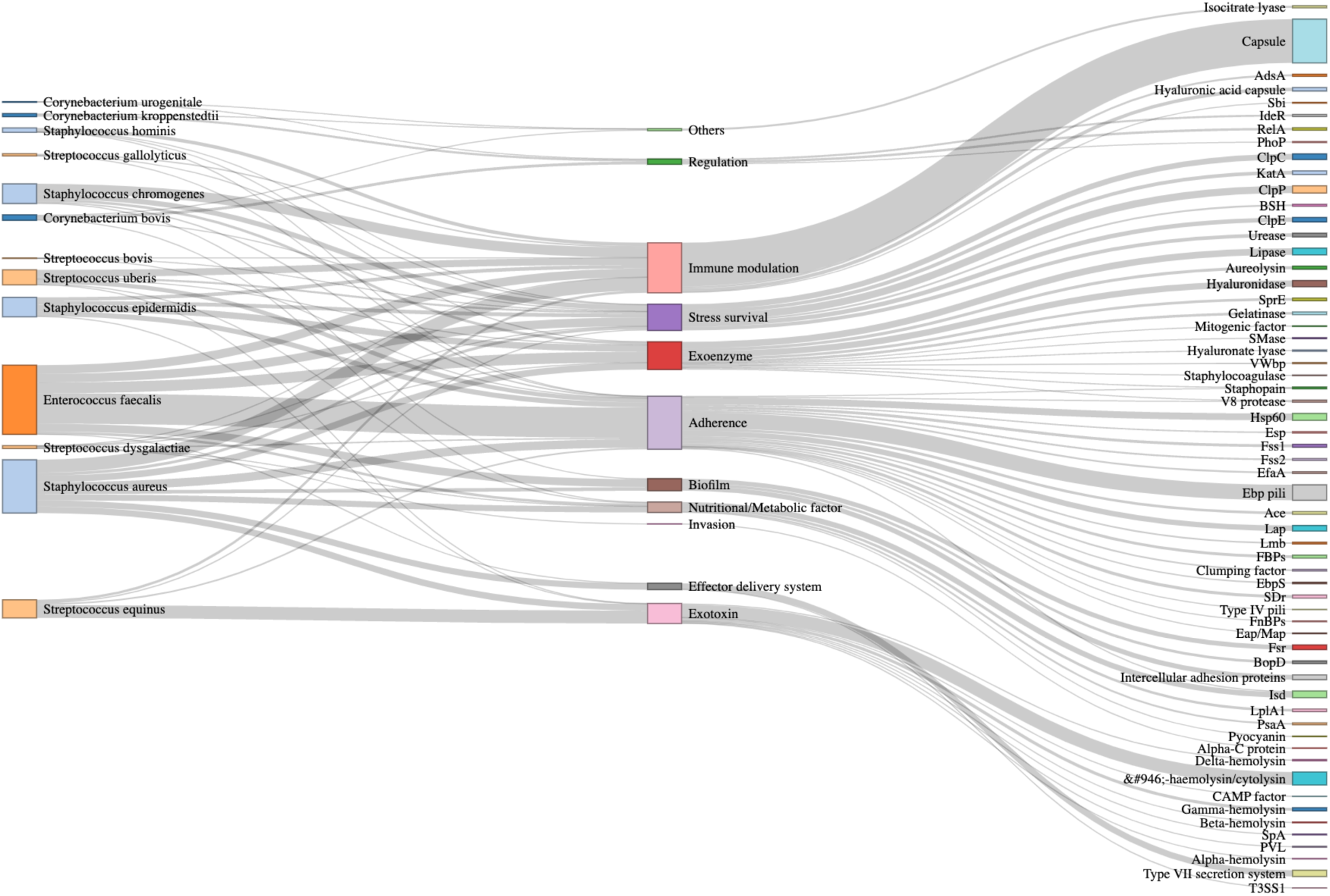
Sankey plot illustrating the relationship between bacterial taxa, virulence factor categories, and specific virulence factors. The thickness of the connecting lines represents the relationships between the taxa, virulence factor categories, and specific virulence factors. Color coding differentiates bacterial taxa and highlights the distribution of virulence traits across the species.

For *Staphylococcus* spp., *S. aureus* showed dominant VFs categories associated to immune modulation (24.53%) and exoenzymes (14.34%), followed by nutritional/metabolic factors (13.58%), exotoxins (12.08%), adherence (11.70%), and effector delivery systems (11.70%). Other categories included biofilm formation (6.79%) and stress survival (5.28%). *S. chromogenes* was predominantly associated with immune modulation (51.02%), followed by stress survival (25.51%) and exoenzymes (13.27%). Adherence (7.14%) and biofilm formation (3.06%) were less frequent. In *S. epidermidis*, exoenzymes were most abundant (36.08%), followed by adherence (22.68%), stress survival (19.59%), and immune modulation (18.56%). Exotoxins accounted for a minor proportion (3.09%). *S. hominis* was heavily skewed towards immune modulation (77.27%), with smaller contributions from stress survival, adherence, and exoenzymes (13.64%, 4.55%, and 4.55%, respectively).

In species of *Streptococcus*, adherence, immune modulation, and stress survival were equally represented (33.33% each) for *S. bovis*. *S. dysgalactiae* demonstrated a focus on adherence (50%), followed by exoenzymes and stress survival (14.29% each), with other categories such as immune modulation, invasion, and nutritional/metabolic factors at 7.14% each. For *S. equinus*, exotoxins dominated (71.11%), with smaller contributions from immune modulation (13.33%), adherence (8.89%), and stress survival (6.67%). *S. gallolyticus* exhibited a strong preference for immune modulation (63.64%), with stress survival and adherence making up 27.27% and 9.09%, respectively. Finally, *S. uberis* showed a varied distribution, with immune modulation (46.67%) and adherence (28%) being the most common categories, followed by stress survival (13.33%), nutritional/metabolic factors (6.67%), exoenzymes (4%), and exotoxins (1.33%).

Since the VFDB database lacks information on members of the *Aerococcaceae* family (as it only includes experimentally verified virulence factors), the prediction of virulence factors for microorganisms associated with this taxon was performed using category V (Defense mechanisms) from the COG database as a reference. This analysis identified six protein-coding genes that may enable bacteria of this taxon to colonize the bovine mammary gland and enhance their pathogenic potential: hrtA (putative hemin import ATP-binding protein), hsdM (site-specific DNA-methyltransferase, adenine-specific), ylbB (efflux ABC transporter, permease protein), yheH (ABC transporter transmembrane region), ywjA (ABC transporter transmembrane region), and XK27_06785 (putative ABC transport system ATP-binding protein, ABC.CD.A).

In the analysis of antimicrobial resistance (AMR) genes, a total of 144 genes were annotated using three databases: ResFinder, CARD, and ARG-ANNOT (**Figure 14**). The top five antibiotic classes represented in the dataset were fluoroquinolones (15%), macrolides (10.2%), tetracyclines (9.6%), aminoglycosides (7.6%), and rifamycins (5.8%).

**Figure 14.**
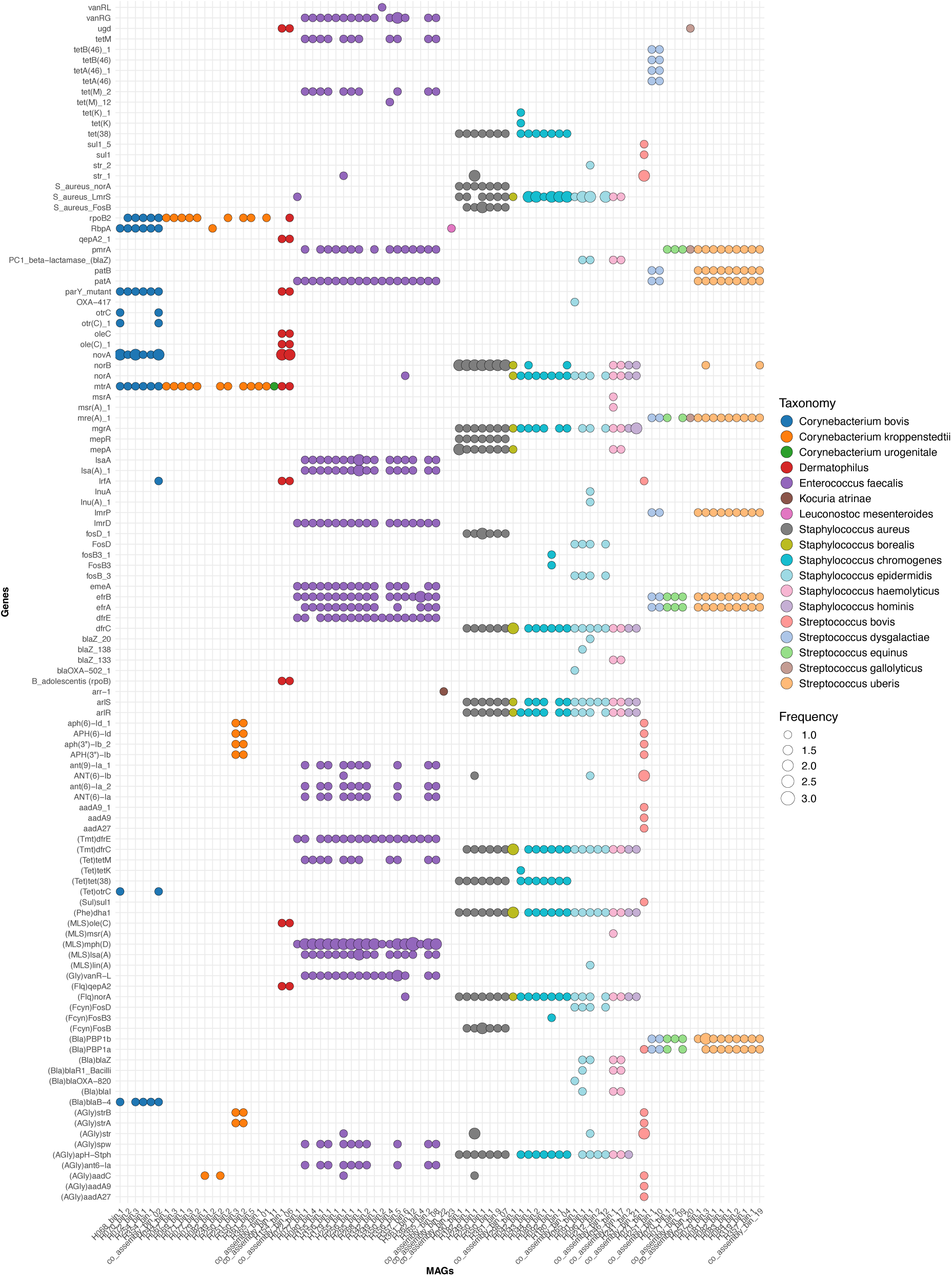
Dot plot showing the distribution of antibiotic resistance genes across metagenome-assembled genomes. Dot sizes are proportional to the frequency of each gene as annotated by three databases: ResFinder, CARD, and ARG-ANNOT. Each dot is color-coded by MAG, highlighting the presence and prevalence of specific ARGs across different MAGs. Larger dots indicate higher consensus across the databases regarding the detection of that gene.

When stratified by bacterial species, distinct patterns of AMR gene distribution were observed. *Corynebacterium* spp. exhibited a high abundance of genes associated with rifamycins (20.4%), macrolides (14.6%), penams (14.0%), aminoglycosides (12.7%), and aminocoumarins (12.1%). For *Staphylococcus* spp., the most abundant AMR genes were associated with fluoroquinolones (18.2%), acridine dyes (11.2%), tetracyclines (10.8%), aminoglycosides (8.3%), and phenicols (7.9%). Similarly, *Streptococcus* spp. displayed a predominance of genes linked to fluoroquinolones (24.2%), macrolides (14.2%), beta-lactams (11.1%), rifamycins (10.4%), and tetracyclines (6.6%). For *E. faecalis*, AMR genes were most frequently associated with macrolides (13.6%), fluoroquinolones (11.3%), lincosamides (8.1%), tetracyclines (8.1%), and aminoglycosides (8.0%). *Dermatophilus* spp. demonstrated a high prevalence of genes for aminocoumarins (22.9%), fluoroquinolones (20%), macrolides (14.3%), ciprofloxacin (8.6%), and oleandomycin (8.6%). In *Kocuria atrinae*, AMR genes were predominantly annotated to rifamycins (35.7%) and fluoroquinolones (28.6%), with additional annotations to aminocoumarins, macrolides, oleandomycin, oxazolidinones, and peptides (each at 7.1%). Lastly, *Leuconostoc mesenteroides* had only one annotated AMR gene, which was associated with rifamycins. These findings underscore the diversity of AMR gene profiles across MAGs reconstructed from metagenomes obtained from milk samples, highlighting species-specific adaptations to antimicrobial pressures.

#### 3.5.3. Prediction of bacteriocins and bioactive compounds

We used BAGEL4 to identify gene clusters in the MAGs DNA involved in the biosynthesis of Ribosomally synthesized and Post translationally modified Peptides (RiPPs) and (unmodified) bacteriocins. In total, 27 gene clusters were predicted (**Figure 15A**). Bacterial species annotated as *S. aureus* the highest number of different classes such as 159.2; Pediocin (only predicted in the MAG H249_bin.1, 100% identity with bacteriocin immunity protein from *Pediococcus pentosaceus*), 185.1;BacCH91, putative_bacteriocin (Lactococcin_972 – PF09683), 314.1; Auto_Inducing_Peptide_I, 491.1; Lantibiotic, 96.1; putative_lantibiotic (BsaA2), and Sactipeptides. Four areas of interest were identified in MAGs assigned to *E. faecalis* (62.3;enterolysin_A, 63.3; Enterolysin_A, Lasso_peptide, and Sactipeptides), *S. chromogenes* (145.1; Subtilosin_A , 314.1;Auto_Inducing_Peptide_I, 315.1;Auto_Inducing_Peptide_II, and Sactipeptides), *S. uberis* (157.1; uberolysin, 163.2; Penocin_A, 42.2;Bovicin_255_peptide, and 61.3;Dysgalacticin), and *C. kroppenstedtii* (189.2;protease-activatedantimicrobialprotein(PAMP), 42.2;Bovicin_255_peptide, 84.1;Nukacin_A_(NukacinISK-1), and 97.2;Enterocin_X_chain_beta). Three gene clusters were identified in *S. epidermidis* (Sactipeptides, 314.1;Auto_Inducing_Peptide_I, and 64.2;Delta-lysinI), *Dermatophilus* spp. (Sactipeptides, 64.1;Microbisporicin_(NAI-107), and 93.3;Zoocin_A), *S. hominis* (Sactipeptides, 314.1;Auto_Inducing_Peptide_I, and 315.1;Auto_Inducing_Peptide_II), and *S. dysgalactiae* (13.1;Butyrivibriocin, 219.1;Streptolysin, and 58.1;Macedocin). Two gene clusters were identified in *S. equinus* (42.2;Bovicin_255_peptide and 84.1;Nukacin_A_(NukacinISK-1)) and *S. haemolyticus* (315.1;Auto_Inducing_Peptide_II and Sactipeptides). Only one gene cluster was predicted for *S. gallolyticus* (LAPs), *S. bovis* (159.2;Pediocin), *S. borealis* (Sactipeptides), and *L. mesenteroides* (97.2;Enterocin_X_chain_beta).

**Figure 15.**
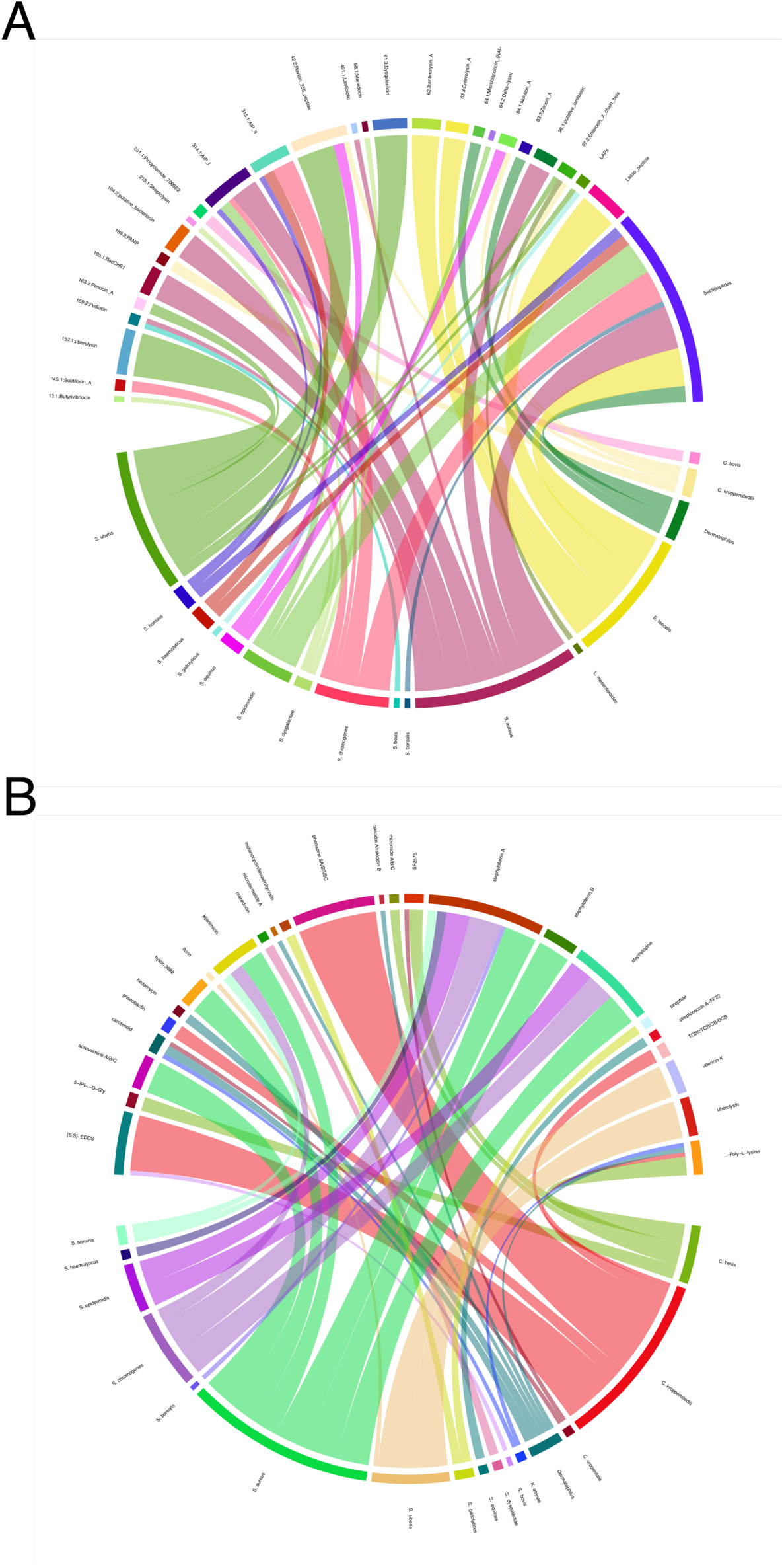
Chord diagram illustrating the relationships between bacteriocin **(A)** gene clusters predicted using BAGEL4 across the different MAGs. Chord diagram depicting the diversity and distribution of secondary metabolite gene clusters identified with antiSMASH **(B)** across the same MAGs. Connections highlight shared features or pathways among the identified clusters, providing insights into the biosynthetic capabilities within each taxon

Genome mining using AntiSMASH identified 322 biosynthetic gene clusters (BGCs) across all analyzed MAGs. These BGCs were classified into 26 distinct types of biosynthetic pathways associated with the production of secondary/specialized metabolites. The ten most abundant pathways were non-ribosomal peptide synthetase (NRPS) (12.7%), terpene (11.1%), non-ribosomal peptide siderophore (NI-siderophore) (10.8%), type III polyketide synthase (T3PKS) (10.2%), ribosomally synthesized and post-translationally modified peptides (RiPP-like) (9.6%), cyclic lactone autoinducer (9.3%), opine-like metallophore (5.4%), NRPS-like (4.8%), aminopolycarboxylic acid (3.9%), and type I polyketide synthase (T1PKS) (3.0%).

When the analysis was stratified by bacterial species and limited to hits with similarity associated with known clusters, several species-specific putative metabolites were identified (**Figure 15B**). In *C. bovis*, the annotated metabolites included Œµ-Poly-L-lysine, 5-isoprenylindole-3-carboxylate β-D-glycosyl ester (here abbreviated as 5-IPI-β-D-Gly), SF2575, and rhizomide A/B/C. For *C. kroppenstedtii*, the identified metabolites were phenazine SA/SB/SC, [S,S]-ethylenediamine-disuccinate ([S,S]-EDDS), trichrysobactin/cyclic trichrysobactin/chrysobactin/dichrysobactin (here abbreviated as TCB/cTCB/CB/DCB), griseobactin, and ε-Poly-L-lysine. *C. urogenitale* showed the presence of SF2575 and carotenoid. Metabolites detected in *Dermatophilus* spp. included carotenoid, hedamycin, microtermolide A, rakicidin A/B, and ε-Poly-L-lysine. The compound ε-Poly-L-lysine was the sole metabolite identified in *K. atrinae*. For *S. bovis*, [S,S]-EDDS was annotated, while macedocin was specific to *S. dysgalactiae*. *S. equinus* displayed streptococcin A-FF22, whereas mutanocyclin/leuvalin/tyrvalin and streptide were annotated for *S. gallolyticus*. In *S. uberis*, the detected putative metabolites included uberolysin, ubericin K, and iturin. *S. aureus* was associated with aureusimine A/B/C, hyicin 3682, staphyloferrin A, staphyloferrin B, staphylopine, and kijanimicin. For *S. borealis*, staphyloferrin A was identified, while *S. chromogenes* yielded staphyloferrin A, staphylopine, and kijanimicin. Staphyloferrin A and staphylopine were predicted in *S. epidermidis*, and *S. haemolyticus* showed staphyloferrin A. Lastly, in *S. hominis*, kijanimicin and staphyloferrin A were annotated. The AntiSMASH analysis revealed a group of bioactive compounds shared among several Staphylococci species, including staphyloferrin A, staphylopine, and kijanimicin. In contrast, unique metabolic signatures were observed in specific species, such as *S. aureus*, *C. kroppenstedtii*, *S. uberis*, and *C. bovis*. These findings suggest a potential common production of certain bioactive compounds, here predominantly identified among Staphylococci, while other microorganisms, such as *C. kroppenstedtii*, appear to exhibit a more species-specific metabolite spectrum.

## 4. Discussion

Bovine mastitis is widely recognized as the most economically significant disease affecting dairy cattle ^66^. Its multifactorial nature poses substantial challenges to effective management through zootechnical and veterinary interventions^67^. In Norway, where the disease is relatively well-controlled and possesses one of the lowest antimicrobial use in livestock in the world, studies on microbial ecology offer valuable insights into potential taxa associated with udder health, contributing to advancements in prevention and management strategies ^68^.

Somatic cell count (SCC) is globally used as a proxy for diagnosing intramammary infections and monitoring mastitis control throughout lactation. Elevated SCC values are typically associated with a recent or ongoing infection, often confirmed by a positive bacterial culture or, more recently, by PCR targeting specific taxa ^69^. The beginning and end of lactation are generally considered critical periods for udder health due to increased susceptibility to infections. In this study, the lactation period significantly influenced SCC levels, with lower counts observed during early and mid-lactation, while higher counts were recorded near the dry-off period in 2022 and late lactation in 2023, which is in agreement with the scientific literature ^70^.

Mastitis-causing pathogens are a primary driver of elevated SCC in the udder, with various bacterial species implicated. These pathogens are conventionally classified into two main groups: major and minor mastitis-causing pathogens ^3^. In our study, major pathogens such as *S. aureus*, *Enterococcus* spp., and environmental streptococci were prominent, whereas minor pathogens like *S. chromogenes*, *S. haemolyticus*, *S. epidermidis*, and *C. bovis* were less frequently reported. Apart from the elevated culturing and detection of *E. faecalis* in milk samples from the mid-lactation period of 2023, our findings align with a recent prevalence study of udder pathogens in Norwegian dairy cows between 2019 and 2020 ^71^. The high occurrence of *E. faecalis* coincides with the period when animals transition from freestalls to pasture. As reported elsewhere, *Enterococcus* is considered both a commensal organism and an opportunistic pathogen, possessing a remarkable ability to survive under adverse conditions. It is commonly found as part of the normal gut microbiota in animals, which may facilitate its dissemination under specific environmental or management practices ^72^.

Bacteriological culturing of milk samples aids in guiding mastitis treatment decisions by identifying the causative bacteria of intramammary infections. However, as not all mastitis-causing bacteria require treatment, this approach, from a microbial ecology perspective, overlooks the presence of a substantial fraction of microorganisms within a given sample due to the fastidious nature of some microbial taxa ^73^. In our study, two groups of samples classified as negative (i.e., absence of mastitis-causing pathogens according to the gold standard) were identified. The first group (n = 72) exhibited high colony counts (30–300 CFUs) but had somatic cell counts (SCC) below 100,000 cells/mL. The second group (n = 57) also showed high colony counts but had a median SCC of 2.9 × 10⁵ cells/mL. Although these microorganisms are not routinely identified, they may contribute to elevated SCC levels. For example, the genus *Lactococcus* has emerged as a potential group of emerging mastitis pathogens, yet it is rarely documented in conventional culturing reports ^74,75^.

In an effort to overcome the limitations of culture-dependent methods and complement them, numerous studies have utilized metataxonomic analysis to explore the microbial composition of bovine milk in both health and disease ^21,76–78^. In this longitudinal study, we conducted a comprehensive investigation of the microbial composition at the quarter level in over 300 milk samples, aiming to uncover microbial succession patterns associated with bovine mastitis. The lower diversity indices observed in samples with high somatic cell counts are a common finding in ecological studies of the bovine mammary gland. This reduction in diversity reflects the dominance of specific microbial taxa, which disrupt the homeostasis of the milk microbial community, resulting in a dysbiotic state within the affected quarter ^19,23,76^.

In this study, we observed that both the composition and relative abundances of bacterial taxa at the phylum and genus levels varied across lactation periods. Specifically, an enrichment of *Proteobacteria* during the early lactation period (EL-2023) was associated with a lower frequency of samples exhibiting high somatic cell counts. This phylum is linked to cases of clinical mastitis, such as those caused by *Escherichia coli*. However, *Proteobacteria* are also frequently observed at high relative abundances in healthy milk samples. In contrast, cases of subclinical mastitis are more commonly associated with the phylum *Firmicutes*, including bacterial species such as *Staphylococcus* spp. ^17^. The increase in the phylum *Actinobacteriota* in the milk microbiota during the mid- and late-lactation periods may be attributed to dietary changes and seasonal factors, as it aligns with rising temperatures and the transition to a pasture-based system. Recent studies suggest that the ruminal microbiota and climatic conditions can significantly impact udder health ^79,80^.

The microbial diversity observed at the genus level in this study aligns with findings from previous research conducted on the same herd and globally, with taxa such as *Staphylococcus*, *Corynebacterium*, *Streptococcus*, *Enterococcus*, *Aerococcus*, *Kocuria* and *Romboutsia* among the most prevalent ones ^28,81,82^. To further investigate the structure and dynamics of the milk microbiome community, we applied a dimensionality reduction method commonly used in human microbiome studies to explore associations with parameters of interest ^83^. This approach provided valuable insights, particularly regarding sample classification, bridging the gap between culturomics and culture-independent methods. By combining these approaches, we increased the detection of specific pathogens in samples initially classified as negative, highlighting the potential of this methodology to enhance microbial detection and classification. Conventional bacteriological examination remains the most widely used method for diagnosing bovine intramammary infections, however, it is estimated that over 30% of milk samples yield negative culture results or fail to identify the causative agent ^84,85^. According to Smistad et al. (2023) ^71^, the proportion of samples classified as culture-negative in subclinical mastitis cases can exceed 20% in Norway, while culture-negative samples from clinical mastitis cases account for approximately 10%. Overall, culture-negative mastitis is often associated with bacterial species present below the detection threshold of culture methods or with fastidious microorganisms ^11,86^.

The application of dimensionality reduction enabled the evaluation of the relative abundance of key bacterial genera within each cluster, offering valuable insights into their prevalence across various samples. *Staphylococcus*, *Corynebacterium*, and *Aerococcus* were consistently distributed among the clusters, bringing out their widespread occurrence and variability in the analyzed milk samples. This observation confirms the significant prevalence of these taxa in the milk microbiome, irrespective of whether the sample originates from a healthy udder or a mastitis case. *Staphylococcus* species stand out for their high contagious potential within herds and can be present in both healthy and mastitic quarters, with their prevalence being highly strain-dependent. Although the specific role of *Corynebacterium* spp. in udder health remains unclear ^87^, it has been hypothesized that natural infections may confer protection against major mastitis pathogens, such as *S. aureus*. This is consistent with our findings and may be explained by the production of antimicrobial compounds or modulation of the udder immune system, as reported in previous studies ^88–90^. Meanwhile, *Aerococcus* has been primarily associated with healthy milk samples, and its occurrence with non-aureus staphylococci has been negatively correlated with *E. coli* in clinical mastitis ^26,91^.

Within the bacterial genera evaluated in this study, the remarkable diversity of *Corynebacterium* and *Staphylococcus* species observed through taxonomic analysis stands out, aligning with findings from the scientific literature. According to Seshadri et al. (2022) ^90^, only 30%–50% of the projected actinobacterial phylogenetic diversity has genomic representation via isolates or metagenome-assembled genomes. This limited representation may account for the high number of amplicon sequence variants annotated as uncultured bacteria within the *Corynebacterium* genus in our study, highlighting the need for further metagenomic research utilizing shotgun sequencing to improve taxonomic resolution. In a study designed to survey and compare the microbial diversity of external and teat canal epithelial microbiomes using different sequencing approaches, Dean et al. (2021)^92^ reported that the most diverse operational taxonomic units across all sampling locations were assigned to the *Corynebacterium* genus. Among these, *C. efficiens*, *C. marinum*, *C. maris*, and *C. xerosis* constituted the majority of species-level identifications (mean relative abundance exceeding 1%). Except for *C. bovis*, many *Corynebacterium* spp. present in milk samples can also be isolated from the housing environment and milking-related niches ^93^, reflecting the ubiquity of these microorganisms in the dairy environment.

The majority of the ASVs annotated within the *Staphylococcus* genus belong to the Non-aureus Staphylococci and Mammaliicocci (NASM) group, one of the most frequently isolated bacterial groups from milk samples ^94^. In this study, NASM species such as *S. borealis*, *S. chromogenes*, *S. hominis*, *S. epidermidis*, *S. haemolyticus*, *S. xylosus*, *S. saprophyticus*, and *S. simulans* were identified across the milk samples. Compared to coagulase-positive staphylococci, these species generally exhibit lower virulence but can occasionally cause mastitis ^95,96^. Moreover, they are often overlooked in routine laboratory diagnostics, as recently reported for *S. borealis* and *S. rostri* ^97^. Nonetheless, NASM has garnered increasing attention due to its genetic diversity and its ability to antagonize colonization by *S. aureus* ^98,99^. This antagonistic effect is attributed to the production of various antibacterial molecules, such as bacteriocins, which may play a critical role in shaping microbial dynamics and udder health ^100^.

We observed that the composition of milk microbiota varied both at the cow level and the quarter level. When an intramammary infection was established by *Staphylococcus*, a persistent dysbiosis was detected in subsequent samples, a phenomenon not observed for *Streptococcus*. While a considerable number of cross-sectional studies have examined the bovine mammary gland ^68,77,101^, there is limited knowledge regarding the dynamics of milk microbiota at the quarter level in longitudinal studies over a complete lactation cycle. Our findings align with the scientific literature, which indicates that once *Staphylococcus* establishes dominance within a quarter, it becomes challenging for the cow to eliminate it ^21^. This aligns with the wide variety of mechanisms deployed by *S. aureus*, such as biofilm formation, the emergence of small colony variants, and its capability to invade professional and nonprofessional cells. These mechanisms protect *S. aureus* from the cow’s innate and adaptive immune responses as well as from antimicrobial treatments ^102^. In contrast, our findings regarding *Streptococcus* are consistent with a recent study by Urrutia-Angulo et a. (2024) ^19^ which reported a microbiome profile suggesting full recovery from dysbiosis caused by *Streptococcus* one and three months after the diagnosis of clinical mastitis. Based on the ASVs identified for the genus *Streptococcus* in this study, there is a high prevalence of sequences attributed to *S. uberis*, a primarily environmental pathogen ^103^. This bacterium can cause either persistent subclinical inflammation or severe inflammation associated with transient mastitis, depending on the strain involved ^104^. These results highlight distinct microbiota dynamics depending on the causative pathogen and underscore that *S. aureus* remains a major concern.

Through shotgun sequencing and genome-centric metagenomics, we aimed to gain deeper insights into the functional potential of the microbial community and to reconstruct genomes to better understand microbial interactions within the udder. Despite progress in this field, a significant gap remains in the scientific literature, which can be addressed through more studies incorporating shotgun sequencing to complement metataxonomic analyses at the quarter level. One major challenge is the contamination of eukaryotic DNA from mammalian cells during metagenomic DNA extraction. Combined with the low microbial biomass typically present in milk samples, this reduces the proportion of microbial reads, resulting in poor resolution of the microbial community. Our research group recently developed a method to address this issue ^23^, although further optimization is required for milk samples with low somatic cell counts.

The microbial diversity observed from the reconstructed microbial genomes in the metagenomic sequencing data included the most prevalent bacterial genera identified through amplicon sequencing (*Staphylococcus*, *Corynebacterium*, *Streptococcus*, *Enterococcus*, *Aerococcus*, *Kocuria*, and *Romboutsia*). Notably, a diverse array of draft genomes was detected, particularly within the genera *Staphylococcus* and *Corynebacterium*. Of particular interest is the high abundance of the MAG *C. kroppenstedtii* HC-IA_H069.1. This species belongs to the *C. kroppenstedtii*-like group, which includes *C. parakroppenstedtii* and *C. pseudokroppenstedtii* ^105^. Species of this group have gained recognition for their involvement in breast abscesses and granulomatous mastitis in women ^106^. However, the role of *C. kroppenstedtii*-like strains in bovine udder health and disease remains unclear and warrants further investigation. Species of the genus *Corynebacterium* are well-known for their ability to produce secondary metabolites and are routinely used as cell factories, exemplified by *C. glutamicum* ^107^. A recent study conducted by Liu et at. (2024) ^108^ reports the discovery of a novel glycolipid involved in the development of granulomatous lobular mastitis. This pathogenic factor called corynekropbactin is secreted by *C. parakroppenstedtii* and may chelate iron, cause the death of mammary cells and other mammary-gland-colonizing bacteria, and increase the levels of inflammatory cytokines. Studies exploring the adaptation of this species to the bovine mammary gland, as well as comparative genomics analyses with human isolates, are needed to better understand its potential implications in the bovine mammary gland.

The MAGs *Facklamia* bacterium HC-CA013, *Kocuria atrinae* HC-CA018, *S. equinus* HC-IA_356.3, *Staphylococcus* bacterium HC-IA_319.2, *S. chromogenes* HC-IA_H042.1, *Peptoniphilus* bacterium HC-CA024, *Corynebacterium* bacterium HC-CA026, and *Corynebacterium* bacterium HC-IA_212.3 were found at higher coverage in milk samples with low somatic cell counts. The genera *Facklamia* and *Kocuria* have been reported as part of the milk microbiota of healthy bovine quarters and human precolostrum ^109,110^. *Facklamia* belongs to the family *Aerococcaceae*, which comprises nine genera; recently, some species of *Facklamia* were reclassified, such as *F. tabacinasalis*, now *Ruoffia tabacinasalis* ^111^. Meanwhile, *Kocuria* is a member of the family *Micrococcaceae*, and the genus was established by subdividing the genus *Micrococcus* ^112^. Metabolically, *Aerococcaceae* are classified as lactic acid bacteria and can produce lactic acid as the primary product of carbohydrate fermentation ^113^, a pathway highlighted in this study by the enrichment of COG category G (Carbohydrate transport and metabolism). Their metabolism reflects adaptation to low-oxygen, nutrient-rich environments, such as milk. Although these microorganisms are well-suited to milk, studies exploring the crosstalk between the identified MAGs and their roles in bovine udder health remain largely unexplored.

A comprehensive understanding of the genetic variation among mastitis-causing pathogens in bovine populations, including temporal trends, is essential for developing effective mastitis control strategies. This is particularly important because the severity of the disease and its impact on milk production are, at least in part, influenced by the genetic and phenotypic characteristics of specific pathogenic microorganisms. In this study, we conducted an *in silico* multilocus sequence typing (MLST) analysis on metagenomic samples and the metagenome-assembled genomes (MAGs) derived from these datasets. Among the lineages identified, all *S. aureus* sequences were associated with the ST133, which has been recognized as a globally distributed sequence type predominantly associated with livestock such as large and small ruminants ^114,115^. Regarding *E. faecalis*, all sequences analyzed in this study were assigned to clonal complex 40 (ST40), the most prevalent clonal type of this species. ST40 is globally distributed, with isolates originating from various sources, including livestock, humans, and environmental reservoirs ^116^. This lineage is of particular significance due to its widespread presence in clinical and non-clinical settings. It is frequently isolated from both commensal and pathogenic contexts, demonstrating its adaptability across different environments. *S. dysgalactiae* ST302 has been previously identified in cows, with evidence suggesting it is well-adapted to this host ^117^. In contrast, limited information is available in the scientific literature regarding *Streptococcus uberis* ST1409. However, it appears closely related to sequence type 1405 and has been reported to be sensitive to penicillin, tetracycline, and erythromycin^118^. The ST100 identified in *S. epidermidis* has been described as having an animal origin and is reported to cause mastitis in both ewes and bovines ^119^. In contrast, limited information is available regarding ST575. However, one study identified an antimicrobial compound known as epifadin, produced by nasal *S. epidermidis* strain IVK83 belonging to the ST575, demonstrating the ability to eliminate *S. aureus* ^120^. For *S. chromogenes*, ST1 has been reported as one of the most predominant sequence types in a study investigating differences in the occurrence of intramammary infections in freshly calved heifers in Sweden ^121^. In contrast, for ST59 and ST104, further research is needed to elucidate their clinical significance, antimicrobial resistance profiles, and associated virulence factors.

To successfully colonize a host, bacteria possess an array of proteins and other components collectively known as virulence factors ^122^, although this terminology for commensal bacteria is not appropriate and should be replaced by a more appropriate term such as “niche factors” ^123^. Interestingly, many of these factors are not exclusively specialized for host-pathogen interactions but play broader roles in adapting to diverse ecological niches and for cooperation in complex biological ecosystems ^124^. The analysis of virulence factors across bacterial taxa in milk samples revealed distinct patterns. According to Tauch & Burkovski (2015) ^125^, many members within the genus *Corynebacterium* developed into rare pathogens “by chance” and most of the species share a more general adhesion mechanism. In our study, regulatory factors such as iron-dependent repressor (IdeR) and PhoP seem to play an important role in niche adaptation. IdeR in *Mycobacterium tuberculosis* and its homolog, the diphtheria toxin repressor (DtxR) in *Corynebacterium diphtheriae*, are crucial iron-dependent transcriptional regulators ^126,127^. IdeR controls iron acquisition, storage, and virulence-related genes in *M. tuberculosis* ^128^, whereas PhoP gene is the response regulator of the PhoPR two-component systems, which plays a crucial role in the virulence of *C. pseudotuberculosis* ^129^. In *E. faecalis*, adherence genes constituted the largest fraction of virulence factors in this study. Key colonization factors included the collagen adhesin Ace, the endocarditis antigen EfaA, and the endocarditis- and biofilm-associated pili (Ebp), all of which are commonly present in *E. faecalis* isolates ^130,131^. The Esp surface protein is more frequently found in isolates from urinary tract infections and bacteremia ^131^. Ebp pili play a major role in adherence to fibrinogen and a minor role in collagen binding, whereas Ace serves as the primary collagen adhesin^132^. Species of *Staphylococcus* employ diverse immune evasion strategies to persist within host organisms. In this study, immune modulation was identified as the predominant category of virulence factors among *Staphylococcus* species, particularly genes involved in capsule formation (*cap8*), *adsA*, and *sbi*. The *cap8* gene cluster, consisting of 16 open reading frames (*cap8A-P*), is responsible for the synthesis of type 8 capsular polysaccharides, with 11 genes directly contributing to its production^133^. Capsular polysaccharides 8, which is predominant among *S. aureus* isolates from humans, can elicit a robust inflammatory response. However, these capsular types are rapidly eliminated compared to capsule-negative *S. aureus*, which are more efficiently internalized by mammary gland epithelial cells and capable of establishing chronic infections in the host ^134^. Other virulence factors, such as staphylococcal protein A (SpA), staphylococcal binder of immunoglobulin (Sbi), and adenosine synthase A (AdsA), also play crucial roles in immune evasion ^135,136^. Sbi, which is both secreted and cell-envelope associated, enhances bacterial survival in human blood and prevents neutrophil-mediated opsonophagocytosis and modulates B-cell activation ^137,138^. Collectively, these mechanisms enable *S. aureus* to evade host immune defenses effectively, facilitating persistent and recurrent infections. *Streptococcus* species possess various virulence factors that contribute to their pathogenicity. Biofilm production is a prevalent trait, observed in over 70% of *Streptococcus* isolates ^139^. *S. agalactiae* strains frequently carry genes like *cyl* genes, *cfa*/*cfb*, and *hylB* ^140^. The lactoferrin-binding protein is a crucial virulence factor found in *S. dysgalactiae*, *S. uberis* and *S. agalactiae*, with differing effects on cell invasion ^141–143^. Notably, *S. agalactiae* strains carrying the *lmb* gene showed enhanced invasion ability and were associated with reduced antibiotic treatment efficacy ^140^.

Many of the identified virulence factors are directly linked to the high recalcitrance associated with antibiotic use in dairy farming. More recently, shotgun metagenomic sequencing has shown to be a valuable tool to quantify the dynamics of AMR transmission at the human–livestock interface ^144^, as well as implications for antimicrobial stewardship in livestock ^145^. Norway stands out for its notably low rates of antibiotic use and bovine mastitis treatment globally, with benzylpenicillin procaine being the first-line therapy for mastitis ^71^. Currently, antimicrobial treatments are primarily reserved for moderate to severe cases of mastitis during lactation and for addressing subclinical mastitis caused by *S. aureus*, *S. dysgalactiae*, *S. uberis*, or *S. agalactiae* at dry-off ^71^. *Corynebacterium* species exhibit widespread antibiotic resistance, particularly to macrolides, lincosamides, and streptogramins ^146,147^. Additionally, resistance to β-lactams, clindamycin, erythromycin, azithromycin, ciprofloxacin, and gentamicin is also common among various *Corynebacterium* species. Most strains remain susceptible to vancomycin, minocycline, and linezolid^148^. Given the high prevalence and diversity of *Corynebacterium* species in milk samples, this may be linked to their common resistance to beta-lactams ^149,150^, potentially contributing to the persistence of these microorganisms within the herd enrolled in this study. Of food safety concern, *S. aureus* has developed resistance to various antibiotics through mechanisms including enzymatic inactivation, target alteration, antibiotic trapping, and efflux pumps ^151^. Exposure to biocides and dyes can lead to overexpression of efflux pump genes like *mepA*, *mdeA*, *norA*, and *norC*, contributing to multidrug resistance ^152^. Staphylococci of animal origin possess numerous resistance genes conferring resistance to multiple antibiotic classes, including fluoroquinolones, tetracyclines, aminoglycosides, and phenicols. These genes are often carried on mobile genetic elements, facilitating their spread among bacterial populations ^153^. Species of *Streptococcus* have also developed resistance to multiple classes of antibiotics in both humans and various animal species, presenting significant challenges for effective treatment ^154,155^. Key resistance mechanisms include genetic modifications, alterations in target sites, and the activity of efflux pumps. ^156^. The dissemination of resistance genes among streptococci is primarily mediated by transposons and integrative conjugative elements ^155^. While streptococci are typically regarded as sensitive to benzylpenicillin and are not routinely tested in Norway, there have been reports of structural modifications in penicillin-binding proteins caused by point mutations in *S. pneumoniae*, *S. agalactiae*, and *S. dysgalactiae* subsp. *equisimilis*. These changes have been linked to reduced susceptibility to β-lactam antibiotics ^157–159^. Macrolide resistance in *Streptococcus* arises through three primary mechanisms: ribosomal modifications (both post- and pre-transcriptional), active antibiotic expulsion via efflux pumps, and target protection ^160^, and studies have frequently linked this resistance to the presence of the *ermB* gene in *S. agalactiae*, *S. dysgalactiae*, and *S. uberis* ^161,162^. Although fluoroquinolone resistance remains low, it is increasing due to genomic mutations and plasmid-encoded genes, while tetracycline resistance is mediated by ribosomal protection genes, with resistance determinants for macrolides and tetracyclines often co-occurring on mobile genetic elements ^163,164^. *E. faecalis* has been identified as a significant pathogen in bovine mastitis and isolates can exhibit high resistance to tetracycline and erythromycin, with resistance rates of 78.6-87.7% and 28.7-79.0%, respectively ^165,166^. The most prevalent resistance genes include *tetK*, *tetL*, and *tetM* for tetracycline, as well as *ermB* and *ermC* for erythromycin, although the latter were not identified in our study ^165,166^.

Bacteriocins produced by LAB play a significant role in shaping microbial communities and maintaining microbiome homeostasis ^167^. In the context of bovine mastitis, LAB isolated from the mammary glands of cows have demonstrated potential as probiotic strains for prevention ^168^. However, it remains debatable whether the use of LAB aligns with the biology of the mammary gland, as both the bacterial strain and the site of application are critical factors for ensuring effective action without triggering a strong pro-inflammatory response ^169^. Given that coagulase-negative staphylococci are more naturally adapted to the mammary gland, they could represent a viable alternative for the prevention or treatment of mastitis. In this study, several gene clusters were identified across different bacterial species, reflecting the diverse range of molecules utilized by microorganisms in the mammary gland for both competition and pathogenesis. *S. aureus* can express lantibiotics like BsaA2, which provides a competitive advantage in community-acquired strains ^170^. Nukacins are lantibiotics produced by *Staphylococci* and show promising potential for controlling bovine mastitis ^171,172^. *Staphylococcus* species also utilize peptide-based quorum sensing systems, such as the accessory gene regulator (*agr*) system, which employs auto-inducing peptides (AIPs) to regulate various cellular processes ^173^. Enterolysin A, a bacteriocin produced by *E. faecalis*, exhibits antimicrobial activity against Gram-positive bacteria, such as *S. aureus* and *Listeria monocytogenes* ^174,175^. Another toxin, the *E. faecali*s cytolysin, is a two-peptide lytic system active against both prokaryotic and eukaryotic, belongs to the lantibiotic family, and requires two post-translationally modified peptides for its activity cells ^176,177^. The expression of the cytolysin is carefully controlled by a quorum-sensing mechanism, potentially balancing its defensive function against predators with maintaining stealth ^177,178^. *S. agalactiae* strains isolated from bovine mastitis cases in Brazil demonstrated antimicrobial activity against other *Streptococcus* species, with the bacteriocin zoocin A identified in their genomes ^179^. Similarly, bacteriocin-producing strains of *Lactobacillus fermentum* and *S. bovis* isolated from raw milk in Thailand exhibited inhibitory effects against *S. dysgalactiae*^180^.

*Staphylococcus* species produce various secondary metabolites that play crucial roles in microbiome interactions and nutrient acquisition. Staphyloferrin A, a highly hydrophilic siderophore, was isolated from *Staphylococcus hyicus* DSM 20459 and found to be active in 37 other staphylococci ^181^. This compound, along with staphyloferrin B, helps scavenge scarce iron in the environment. Another metallophore, staphylopine, is produced by staphylococci to acquire transition metal ions ^182^. Siderophores play a pivotal role in iron acquisition for bacterial pathogens associated with bovine mastitis, enabling their survival in iron-restricted environments. For example, *S. aureus* can utilize a variety of exogenous iron sources, including siderophores such as ferrichrome and desferrioxamine, as well as heme-containing proteins ^183^. The ability to acquire iron through these mechanisms significantly enhances the competitive fitness of *Staphylococcus* species across diverse ecological niches, contributing to their adaptability and pathogenic potential. A genome-wide CRISPR interference sequencing (CRISPRi-seq) screen carried out by Mårli et al. (2024) ^184^ identified key fitness determinants essential for *S. aureus* growth and survival in milk. The study revealed several genes with differential fitness, highlighting specific adaptations for thriving in this environment, with metal acquisition emerging as particularly critical. In this study, ε-Poly-l-lysine, a bioactive secondary metabolite with potent antimicrobial properties, was identified across various *Corynebacterium* species. This compound is widely utilized in several Asian countries as a food preservative and is also implicated in shaping the microbial dynamics of cheese and skin ecosystems ^185^. The indole-type compound 5-isoprenylindole-3-carboxylate β-D-glycosyl ester has its bioactivity exploited in developing new drugs and it is widely found in actinobacteria ^186^. The compound [S,S]-ethylenediamine-disuccinate ([S,S]-EDDS) was predicted in *C. kroppenstedtii*. According to Costa et a. 2008^187^ , metal complexes of the biodegradable ligand EDDS exhibit antimicrobial activity against fungi and bacteria. Another two noteworthy compounds predicted for *C. kroppenstedtii* using AntiSMASH analysis are a triscatecholamide siderophore and a catechol-peptide structure named griseobactin. These compounds may play a crucial role in iron acquisition under low-iron conditions in the bovine mammary gland, where iron availability varies depending on udder health, lactation stage, and daily milk production ^188,189^. Phenazines, versatile secondary metabolites predicted for *C. kroppenstedtii*, are commonly produced by various bacteria, including *Pseudomonas*, *Burkholderia*, and *Streptomyces* species ^190,191^. These compounds exhibit broad-spectrum antibiotic activity and play a significant role in enhancing the ecological fitness and pathogenic potential of the producing strains ^192,193^. Overall, understanding the potential production of bacteriocins and bioactive compounds may lead to novel anti-infective strategies and microbiome modulation approaches to tackle bovine mastitis.

## 5. Conclusion

Bovine mastitis continues to present a significant challenge to global dairy production, highlighting the need for innovative strategies to manage this complex, multi-etiological disease. This longitudinal study offers valuable insights into the udder microbiome dynamics across lactation stages using both metataxonomic and shotgun metagenomic approaches. Our results reveal substantial variations in somatic cell count and microbiota composition, with dominant species identified in high SCC samples. Through dimensionality reduction, we were able to identify health-promoting bacteria and examine microbial dynamics at the quarter level, both before and after pathogen establishment, demonstrating that *Staphylococcus* remains a primary concern in mastitis pathogenesis. Metagenomic analyses provided a deeper understanding by uncovering pathogen-specific metabolic pathways and reconstructing 142 metagenome-assembled genomes. These findings offer crucial insights into pathogen adaptation, virulence, and resistance mechanisms. The protective roles of *Aerococcus*, *Kocuria*, and Coryneform bacteria warrant further investigation as potential health-promoting bacteria. Additionally, the identification of bacteriocin and biosynthetic gene clusters opens potential pathways for the development of targeted antimicrobial therapies. In conclusion, this study enhances our understanding of the bovine udder microbiome and its role in mastitis pathophysiology. These insights provide a foundation for microbiome-informed strategies that can improve the health and productivity of dairy herds.

## Acknowledgments

This work received economic support from the Norwegian Research Council (grant number 314733), the Faculty of Chemistry. Biotechnology. and Food Science at the Norwegian University of Life Sciences.

## Declaration of generative AI in scientific writing

No generative AI has been used in this article.

