## Supplementary figures and images for "Longitudinal study of the udder microbiome of Norwegian Red dairy cows using metataxonomic and shotgun metagenomic approaches: Insights into pathogen-driven microbial adaptation and succession"

### Supplementary

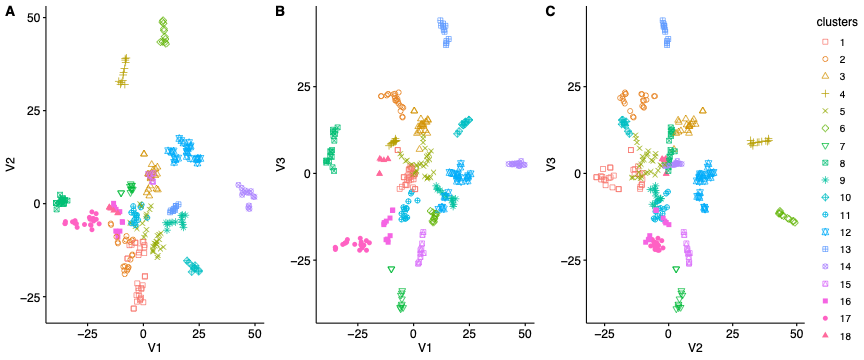

**Supplementary Figure 1.**

**Supplementary Figure 2.**

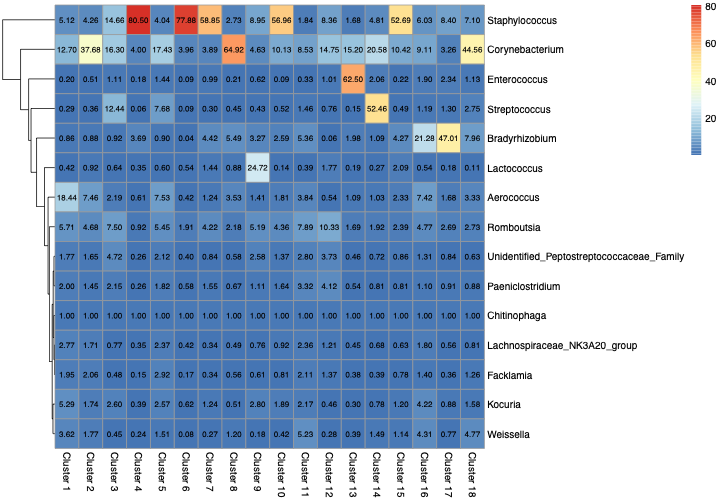
